# Carbonic anhydrase inhibition rescues heart function in the Tg2576 Alzheimer’s mouse model by ameliorating cardiac amyloid-β pathology and neurotrophic-signaling impairment

**DOI:** 10.64898/2026.10.08.757731

**Authors:** Andrea Elia, Rafael Vazquez-Torres, Ashley Carey, Madison Wolf, Varenya R Gade, Micaelly Alves, Silvia Fossati

## Abstract

FDA-approved carbonic anhydrase inhibitors (CAIs) have shown robust efficacy in reducing amyloid-β (Aβ) pathology, neurodegeneration, and cerebrovascular dysfunction in Alzheimer’s disease (AD) and cerebral amyloid angiopathy (CAA) models, highlighting carbonic anhydrase (CAs) as a promising, previously untapped therapeutic target for AD and vascular contribution to dementia. We have recently shown that Aβ aggregates disrupt neuro-signaling and deplete neurotrophic factors (particularly BDNF) in the heart of AD mouse models and human subjects. In this study, we present the first in-depth examination of CAI therapy (acetazolamide-ATZ) in Tg2576 mice, a well-established model of AD and CAA. We confirm that progressive cardiac fibrosis and Aβ deposition in 13-month-old mice severely impair the cardiac neuro-signaling system, marked by a drastic decrease in BDNF and reductions in adrenergic fibers and regenerating nerve endings (TH and GAP-43 markers), leading to significant declines in ejection fraction and fractional shortening compared to age-matched WT littermates. Critically, this study demonstrates that long-term ATZ treatment in Tg2576 mice significantly reduces cardiac Aβ burden, prevents adverse neuro-signaling remodeling, and restores heart physiological function, as assessed via speckle tracking strain echocardiography. These *in vivo* findings were corroborated in human cardiomyocytes, in which CA inhibition reduced human Aβ40 oligomer-dependent disruption of CREB signaling, thus preserving BDNF expression. Together, these results reveal a previously unrecognized cardio-neuroprotective role for CAIs in preventing Aβ-induced cardiovascular remodeling and neuro-signaling disruption in the Tg2576-AD model. Given the established benefits of CAIs on cerebrovascular function and cognition, our findings further support advancing repurposing efforts into clinical trials. CAIs such as ATZ could offer a dual-action therapy approach, addressing both the central nervous system and cardiac pathologies associated with AD and, potentially, the systemic effects of amyloidosis.

**Graphical Abstract:** 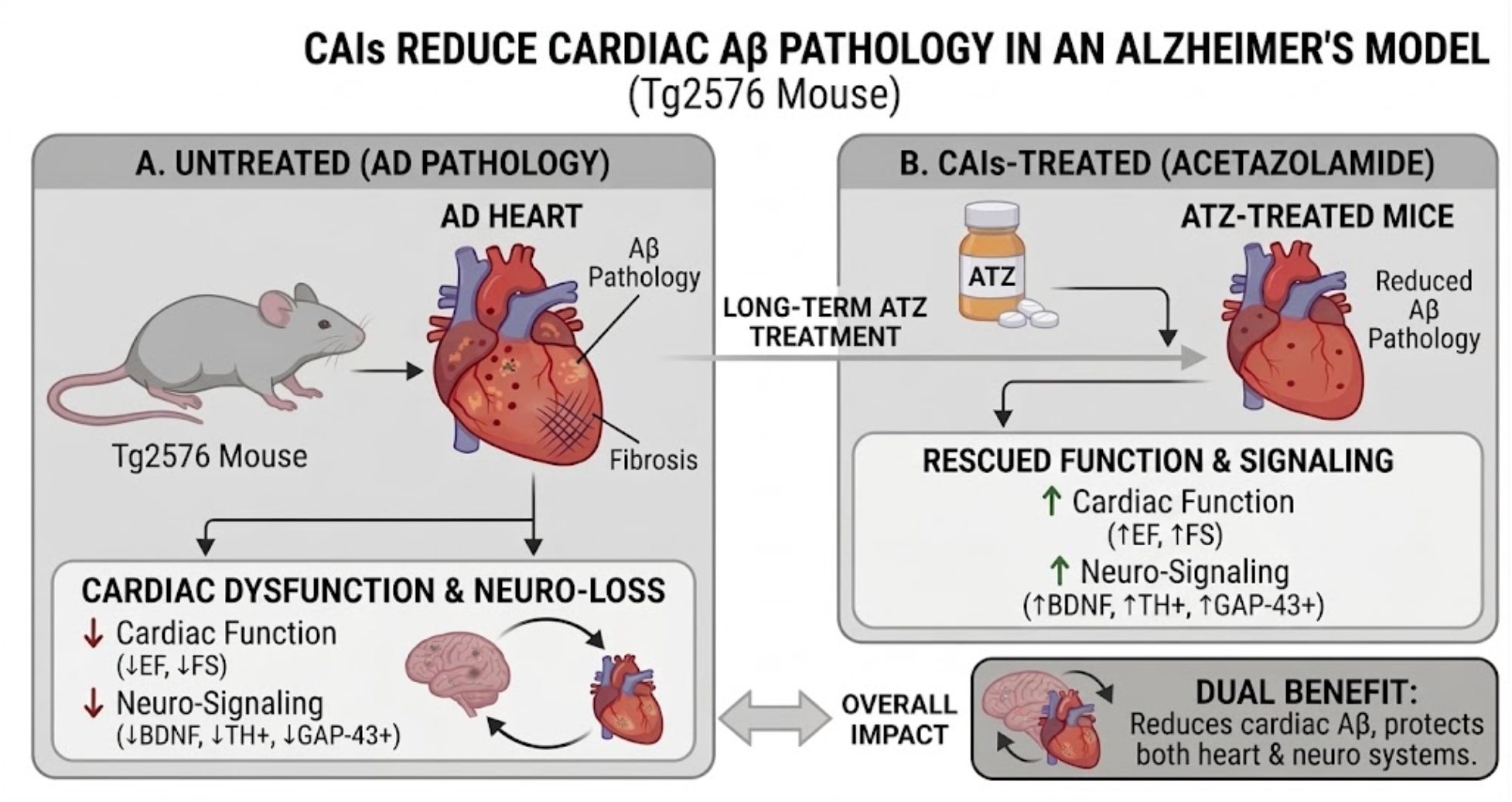

**Effect of Carbonic Anhydrase Inhibition on Cardiac Aβ Pathology in the Tg2576 Heart.** Long-term acetazolamide (ATZ) treatment reduces Aβ deposition and fibrosis in Tg2576-AD mouse hearts, preserving cardiac function (↑EF, ↑FS) and the cardiac neuro-signaling pathway (↑BDNF, ↑TH+, ↑GAP-43+), thereby promoting cardiomyocyte homeostasis and long-term cardiac resilience.

## 1. Introduction

Alzheimer’s disease (AD) represents an urgent and growing global health crisis, with its prevalence projected to triple by 2050, primarily driven by the increasingly aging population (Hebert, Weuve et al. 2013). This multifactorial neurodegenerative disorder is pathologically defined by the accumulation of insoluble amyloid-β (Aβ) plaques within the brain parenchyma and neurofibrillary tangles composed of hyperphosphorylated tau protein. In addition to these hallmark features, approximately 90% of AD cases also exhibit cerebral amyloid angiopathy (CAA), the deposition of Aβ aggregates along cerebral vessel walls, which significantly contributes to disease progression. CAA is increasingly recognized for its role in exacerbating neuroinflammation, impairing neurovascular function, increasing blood-brain barrier (BBB) permeability, reducing brain clearance mechanisms, and accelerating neurodegeneration (Masters, Bateman et al. 2015, Greenberg, Bacskai et al. 2020, Kelly, Brown et al. 2023). Although AD is primarily characterized by central nervous system (CNS) dysfunction, accumulating evidence indicates that its pathological impact extends beyond the brain (Nanclares, Colmena et al. 2025, Elia, Parodi-Rullan et al. 2026, Jiang, O’Bryant et al. 2026). Recent experimental and clinical studies have revealed a compelling association between AD and cardiovascular conditions, including heart failure, ischemic heart disease, and atrial fibrillation (Choubey, Bansal et al. 2023, Sun, Zhuo et al. 2023, Wang, Zhou et al. 2024, Fahim, Yao et al. 2025, Elia, Parodi-Rullan et al. 2026). These findings suggest that AD may represent a systemic disorder with both neurological and extracerebral manifestations, particularly affecting cardiac structure and function (Troncone, Luciani et al. 2016, Sanna, Nusdeo et al. 2019, Elia, Parodi-Rullan et al. 2026). Protein misfolding, a hallmark of AD, is increasingly implicated in cardiac pathology. Recent studies have identified Aβ1-40 and Aβ1-42 aggregates, structurally analogous to those in the brain, and big tau accumulation within the hearts of patients affected by AD, heart failure and dilated cardiomyopathy (Troncone, Luciani et al. 2016, Luciani, Montalbano et al. 2023, Elia, Parodi-Rullan et al. 2026). These aggregates, are associated with diastolic dysfunction (Troncone, Luciani et al. 2016, Aishwarya, Abdullah et al. 2024), myocardial thickening (Jin, Bu et al. 2017, Tublin, Adelstein et al. 2019, Elia, Parodi-Rullan et al. 2026), and electrocardiographic abnormalities (Sanna, Nusdeo et al. 2019, Elia, Parodi-Rullan et al. 2026, Sozio, Sahai et al. 2026), even in AD patients without overt cardiovascular disease. Furthermore, AD and cardiac disorders share overlapping risk factors and genetic profiles (Kivipelto, Ngandu et al. 2005, Gianni, Li et al. 2010). The presence of amyloid deposits in cardiomyocytes and interstitial spaces suggests that Aβ-mediated toxicity may directly impair cardiac structure and function (Willis and Patterson 2013, Troncone, Luciani et al. 2016, Elia, Parodi-Rullan et al. 2026), underscoring a potentially underrecognized peripheral component of AD pathology. Brain Aβ accumulation is known to disrupt neurotrophic signaling pathways, notably reducing the expression of the two main neurotrophic factors (NTFs), nerve growth factor (NGF) and brain-derived neurotrophic factor (BDNF), key regulators of neuronal survival and cognitive function (Allen, Watson et al. 2011). Beyond the brain, BDNF signaling via its receptor, tropomyosin receptor kinase B (TrkB), and downstream effector cAMP response element-binding protein (CREB), also plays essential roles in peripheral tissues, including endothelial cells, muscle cells, and cardiomyocytes, where it supports development, survival, and function (Pius-Sadowska and Machalinski 2017, Kermani and Hempstead 2019, Miranda, Morici et al. 2019, Di Rosa, Zimbone et al. 2021, Elia, Cannavo et al. 2021, Hang, Zhu et al. 2021, Li, Guo et al. 2022). We have recently shown that the progressive neurotrophic decline observed in AD extends to the heart, compromising cardiac innervation and contributing to cardiac dysfunction (Elia and Fossati 2023, Elia, Parodi-Rullan et al. 2026). Decreased circulating BDNF levels have been associated with worsening cardiovascular outcomes, particularly in ischemic heart disease (Feng, Huke et al. 2015, Agrimi, Spalletti et al. 2019, Cannavo, Jun et al. 2023). Accordingly, there is an urgent need to develop innovative therapeutic strategies aimed at limiting Aβ-mediated cardiovascular dysfunction to improve both clinical outcomes and quality of life in patients affected by AD and CAA. Our laboratory has identified carbonic anhydrases (CAs) as key modulators of Aβ-induced toxicity, demonstrating that CA inhibition can prevent mitochondrial-mediated apoptosis in cerebral vascular and neural cells, rescue cognition and ameliorate AD pathology in mouse models of cerebral amyloidosis and AD (Fossati, Giannoni et al. 2016, Solesio, Peixoto et al. 2018, Lemon, Canepa et al. 2021, Anzovino, Canepa et al. 2023, Canepa, Parodi-Rullan et al. 2023, Gutierrez-Jimenez, Rasmussen et al. 2025, Lemon, Canepa et al. 2026). CAs, which regulate pH balance, ion transport, and metabolic processes (Zamanova, Shabana et al. 2019, Mishra, Dunkerly-Eyring et al. 2020), are increasingly recognized as therapeutic targets in both central and peripheral disorders (Huang, Yang et al. 2010, Aggarwal, Kondeti et al. 2013, Swenson 2014, Supuran 2018, Zamanova, Shabana et al. 2019, Mishra, Tiwari et al. 2020, Stoner, Harris et al. 2022) (Lemon, Canepa et al. 2021, Lemon, Canepa et al. 2026). This family of zinc metalloenzymes, found in all vertebrates and located in various intracellular compartments, catalyzes the reversible hydration of CO_2_ and water into bicarbonate and protons (Supuran 2018, Mishra, Tiwari et al. 2020). *In vivo*, the FDA-approved CA inhibitors (CAIs) methazolamide (MTZ) and acetazolamide (ATZ) protected against cerebrovascular damage, neuroinflammation, and rescued cognitive decline in an AD/CAA mouse model (Canepa, Parodi-Rullan et al. 2023). Recent sex-and APOE-specific transcriptomic drug repurposing identified ATZ as one of the 3 best candidates for AD therapy in humans (Vitali, Raikes et al. 2026). Beyond the CNS, elevated CA expression is observed in various cardiac diseases (Vargas and Alvarez 2012, Alvarez, Quon et al. 2013, Torella, Ellison et al. 2014, Zamanova, Shabana et al. 2019), and CAIs have shown cardioprotective effects, reducing blood pressure (Horita, Yakabe et al. 2006, Swenson 2014, Hoff, Zou et al. 2020, Garcia-Llorca, Carta et al. 2024), improving cardiac function, reducing glucose levels, and enhancing diuretic response in heart failure (Alvarez, Johnson et al. 2007, Vargas, Pinilla et al. 2016, Chen, Li et al. 2022, Mullens, Dauw et al. 2022, Martens, Dauw et al. 2023). Despite these promising findings, the role of CA inhibition in mitigating the cardiac effects of amyloidosis and AD remains uninvestigated, highlighting a critical gap and therapeutic opportunity in AD research.

In this study, we investigated, for the first time in the AD/CAA context, the therapeutic effects of CAI treatment on cardiac responses in Tg2576 mice, a well-established model of cerebral amyloidosis (Hsiao, Chapman et al. 1996). These mice develop age-dependent vascular Aβ accumulation in the brain parenchyma, with extensive cerebrovascular Aβ deposits and BBB impairment, closely mimicking human AD/CAA pathology (Robbins, Betensky et al. 2006). We evaluated whether long-term ATZ treatment could attenuate peripheral Aβ burden, with a particular focus on cardiac pathophysiology and neurotrophic signaling dysregulation. For the *in vivo* investigations, we used ATZ treatment due to its favorable pharmacokinetic profile, extensive FDA-approved clinical use, and established dosing protocols (Van Berkel and Elefritz 2018, Gao, Wang et al. 2021, Abutaleb, Elhassanny et al. 2022, Naqvi, Wall et al. 2025). Based on our previous studies demonstrating the beneficial effects of ATZ and MTZ against Aβ toxicity in neuronal and vascular cells, we further tested both compounds in complementary *in vitro* cardiac cell models.

Collectively, this work uncovers a novel therapeutic role for CAIs in mitigating AD-related cardiac and neurotrophic dysfunction and positions these FDA-approved agents as promising candidates for targeting not only the cerebral, but also the cardiovascular manifestations of Alzheimer’s pathology.

## 2. Results

### 2.1. ATZ rescues amyloid-induced myocardial remodeling and preserves cardiac function in Tg2576-AD mice

To evaluate the impact of carbonic anhydrase inhibition on heart amyloid pathology, cardiac architecture and function, we tracked wild-type (WT) and Tg2576 mice over a 13-month timeline (Fig. 1A). At approximately 6 months of age—before the onset of detectable cerebral parenchymal Aβ deposition in this model, which becomes evident by 8 months of age— as demonstrated in (Elia, Parodi-Rullan et al. 2026), Tg2576 mice were randomly assigned to the following groups: Tg2576 untreated or Tg2576+ATZ. ATZ was administered through the diet (20 mg/kg/day) continuously until the end of the study period (13-months of age). Serial echocardiographic assessments until 8 months displayed a homogeneous distribution across all experimental groups (Supplemental Fig. 5A–D), confirming that neither genotype, chronological aging from 4 to 8 months, nor ATZ intervention drove cardiac divergence in younger animals. However, we confirmed that 13-month-old Tg2576 mice exhibited profound systolic impairment, characterized by a significant depression in ejection fraction (EF%) and fractional shortening (FS%) compared to WT controls (Fig. 1B-C, Table 1 and Supplementary Video 1). This aligns with literature evidencing Aβ accumulation and CAA in the Tg2576 brain at this time point (Kawarabayashi, Younkin et al. 2001), and with our previous studies detailing cardiac pathology in this AD model (Elia, Parodi-Rullan et al. 2026). This functional cardiac deterioration was accompanied by severe adverse chamber remodeling, evidenced by sizably elevated left ventricular end-systolic diameter (LVESD) and volume (LVESV) (Table 1). Importantly, ATZ, given in the diet from 6 to 13 months of age, completely rescued cardiac performance, restoring EF% and FS%, while simultaneously attenuating adverse chamber dilation (Fig. 1B-C). As previously demonstrated in this model (Elia, Parodi-Rullan et al. 2026), no detectable changes in diastolic parameters, as evaluated by conventional echocardiography coupled with pulsed-wave Doppler, were observed between experimental groups. Additionally, to further characterize subtle myocardial mechanical changes, we performed high-resolution speckle-tracking echocardiography (Supplemental Fig. 4 and Supplemental Table 1). Tg2576 mice demonstrated a distinct regional impairment in global peak longitudinal strain (Pk%) and longitudinal strain rate (Pk1/s), mainly within the posterior apical and anterior apical segments (Supplemental Fig. 4C-D). This suggests localized sub-endomyocardial deformation that precedes the global dramatic failure onset. In line with the conventional echocardiography results, ATZ treatment significantly reversed these regional and global mechanical impairments, preserving longitudinal structural dynamics (Supplemental Table 2). These functional improvements were corroborated by complementary histological and molecular analyses. Tg2576 hearts exhibited pathological hypertrophic remodeling, severe interstitial fibrosis (Fig 1D), upregulated apoptotic signaling (Fig 1E), increased heart weight to body weight (HW/BW) ratio and a significant enlargement of cardiomyocyte cross-sectional area via Wheat Germ Agglutinin (WGA) staining in the Tg2576 mouse group (Supplemental Fig. 3A-C), as in (Elia, Parodi-Rullan et al. 2026). This hypertrophic response was further confirmed by a robust upregulation of the key hallmark cardiac distress marker, brain natriuretic peptide (BNP) (Geske, McKie et al. 2013, Yoo 2014, Matthia, Setteducato et al. 2022) (Supplemental Fig. 3D-E). Activation of apoptotic pathways in Tg2576 hearts was evidenced by a significant enhancement in cleaved caspase-3 (Cl-Casp3) expression (Fig. 1E-G).

**Figure 1.**
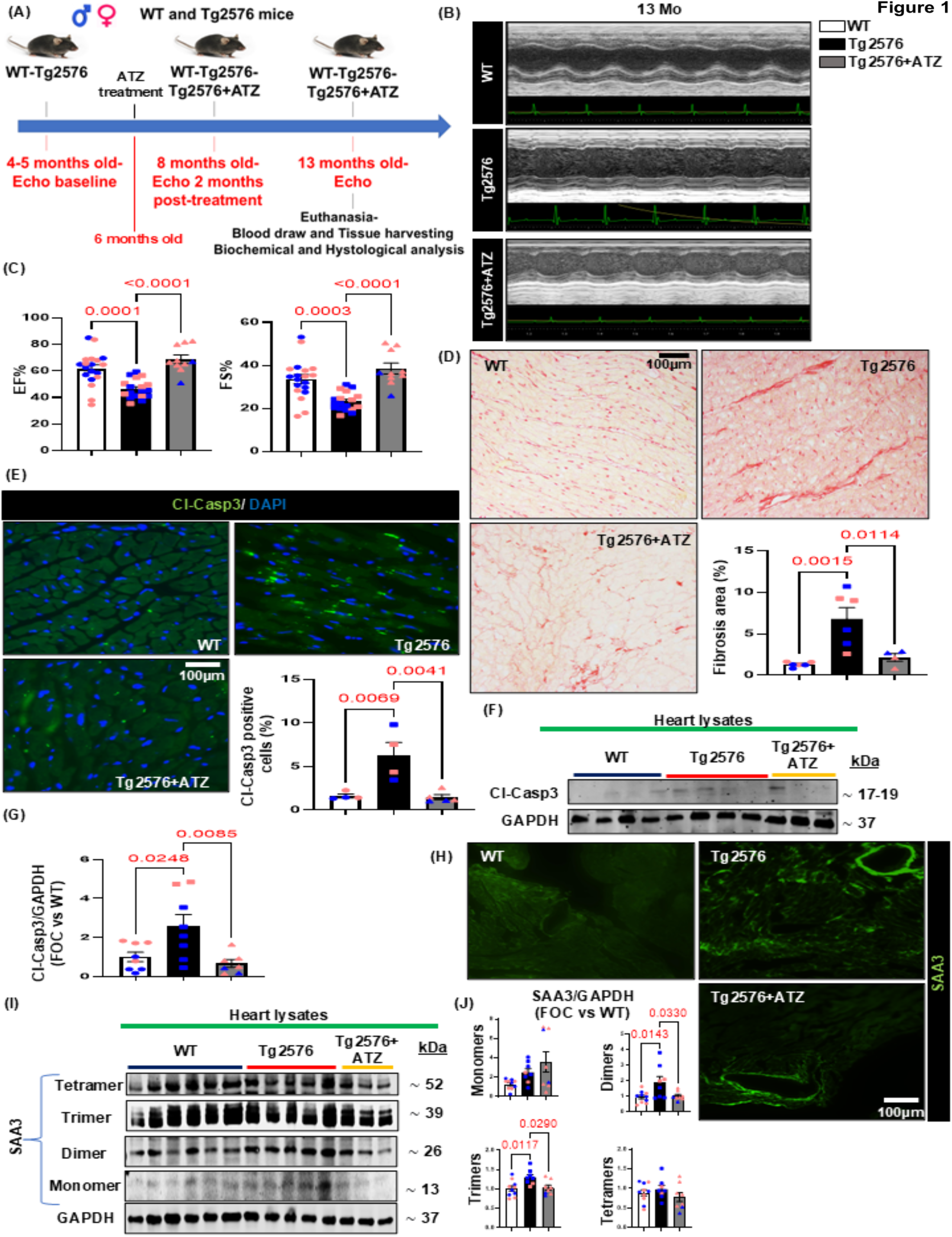
Acetazolamide (ATZ) treatment rescues cardiac dysfunction, suppresses interstitial fibrosis, and attenuates apoptotic signaling in Tg2576 mice. (A) Experimental timeline outlining echocardiographic (Echo) monitoring and ATZ treatment in male and female WT and Tg2576 mice from baseline (4–5 months) through euthanasia (13 months). (B–C) Representative M-mode echocardiographic tracings (B) and corresponding quantification of cardiac functional parameters (C), demonstrating a significant rescue of myocardial contractility in Tg2576 mice in the presence of ATZ treatment. (WT n=19; Tg2576 n=18; Tg2576+ATZ n=10). (D) Representative images of Picrosirius Red-stained myocardial sections (upper panels) and quantitative analysis of fibrotic area (lower panels), showing a marked reduction in interstitial fibrosis in ATZ-treated Tg2576 mice (scale bar: 100 µm) (WT n=6; Tg2576 n=6; Tg2576+ATZ n=4). (E) Representative immunofluorescence images of cleaved Caspase-3 (Cl-Casp3, green) counterstained with DAPI (blue; upper panels) and corresponding quantification of apoptotic cells (lower panels), indicating a robust suppression of myocardial apoptosis by ATZ (scale bar:100 µm) (WT n=4; Tg2576 n=4; Tg2576+ATZ n=5). (F–G) Western blot analysis (F) and relative densitometric quantification (G) of active (cleaved) caspase 3 within the myocardium across experimental cohorts (WT n=8; Tg2576 n=8; Tg2576+ATZ n=7), shows robust reduction of caspase 3 activation by ATZ. (H–J) Representative immunofluorescence images of Serum Amyloid A3 (SAA3, green) deposition (H) and corresponding Western blot quantification (I-J) in cardiac tissues, illustrating that ATZ effectively dampens amyloid-associated SAA3 dimers and trimers upregulation (scale bar: 100 µm) (WT n=10; Tg2576 n=8; Tg2576+ATZ n=8). Data are presented as mean±SEM; individual data points represent individual male (blue) and female (pink) mice. Exact p-values are indicated above the respective comparisons, determined using a multi-group One-way ANOVA with Tukey’s post hoc test.

**Table 1.**
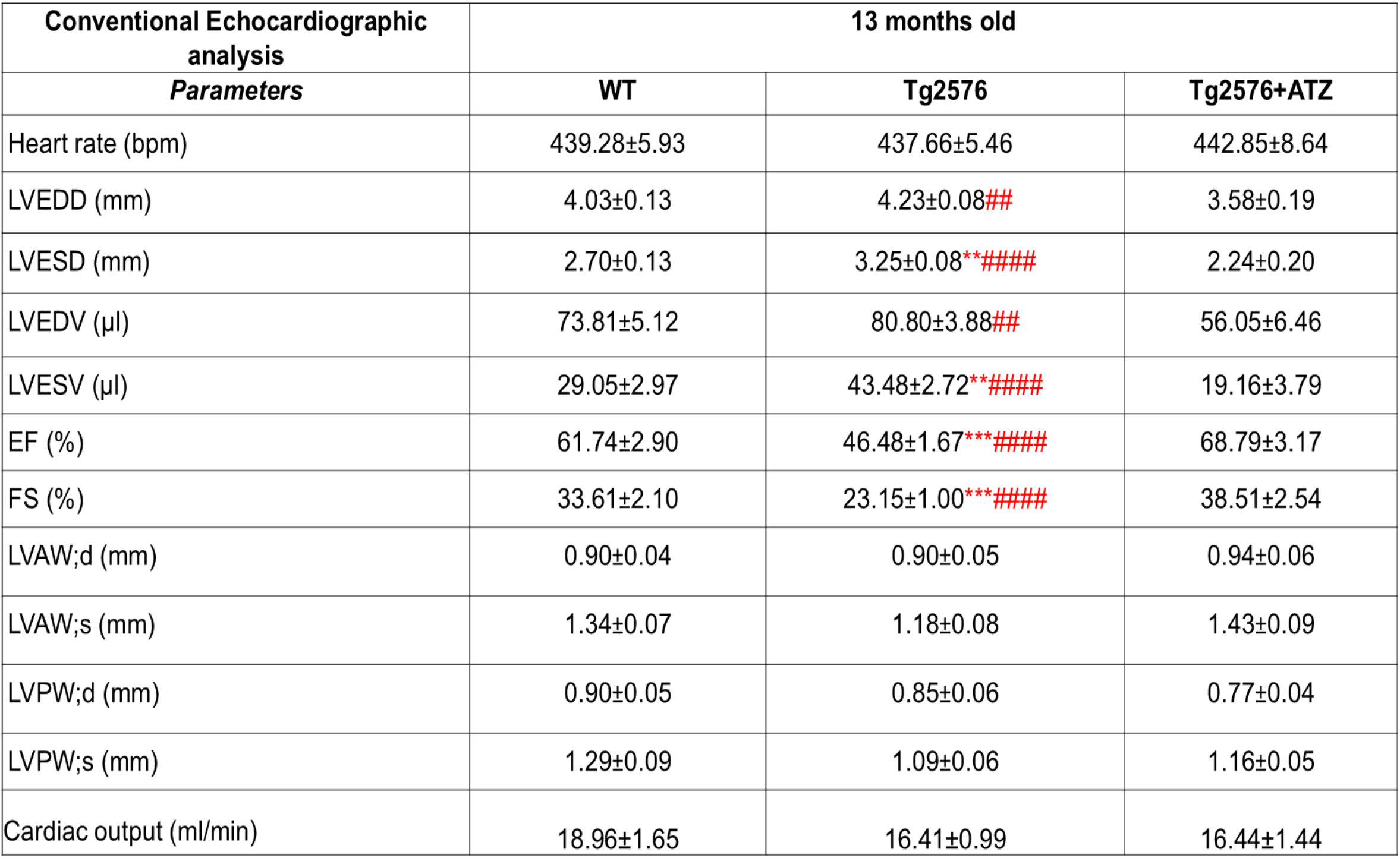
Echocardiographic evaluation of systolic function in WT, Tg2576, and ATZ-treated Tg2576 mice. Conventional echocardiographic parameters were assessed in 13-month-old WT, Tg2576, and Tg2576+ATZ mice: heart rate, left ventricular (LV) end diastolic diameter (LVEDD), end systolic diameter (LVESD), end-diastolic volume (LVEDV), end systolic volume (LVESV), ejection fraction (EF), fractional shortening (FS), anterior wall in diastole (LVAW; d) and systole (LVAW; s), posterior wall in diastole (LVPW; d) and systole (LVPW; s), and cardiac output. n=11-20 mice/group. *P<0.05, **p<0.01, and ***p<0.001 vs WT. P<0.05, ^##^p<0.01, and ####p<0.0001 vs Tg2576+ATZ. (WT n=19; Tg2576 n=18; Tg2576+ATZ n=10). Data are presented as mean ± SEM. One-way ANOVA with Tukey’s multiple comparisons post hoc test was performed between the groups.

Notably, ATZ therapy efficiently hampered all these cardiac AD-driven pathological phenotypes. Specifically, ATZ treatment normalized the HW/BW ratio, limited cardiomyocyte hypertrophy, reduced BNP expression, remarkably decreased interstitial fibrosis (Fig. 1D), and robustly diminished Cl-Casp3-mediated apoptotic cardiac cell death (Fig. 1E-G). We next explored the cardiac accumulation and oligomerization profile of Serum Amyloid A3 (SAA3), a major inflammatory and amyloidogenic mediator implicated in cardiovascular injury and atherosclerotic progression (Morrow, Rifai et al. 2000, Johnson, Kip et al. 2004, Chait, Wang et al. 2021, Cimini, Hansmann et al. 2025), whose pathological signaling cascade may be exacerbated by the progressive deposition of Aβ within the myocardium (Elia, Parodi-Rullan et al. 2026). Immunofluorescence analysis confirmed an increase in SAA3 vascular deposition in 13-month-old Tg2576 hearts compared to WT controls (Fig. 1H). These findings were further corroborated by immunoblot analyses of cardiac lysates, indicating a significantly augmented deposition of SAA3 toxic species spanning dimers and trimers within the Tg2576 myocardium (Fig. 1I-J). Critically, ATZ intervention robustly reduced vascular SAA3 accumulation and the levels of toxic SAA3 dimers and trimers within the transgenic hearts (Fig. 1I-J).

Taken together, these findings strongly demonstrate that ATZ effectively counteracts Aβ-driven cardiac maladaptive remodeling by suppressing fibrosis and hypertrophy, apoptotic signaling cascades, and preserving overall myocardial contractile performance in an AD/CAA mouse model.

### 2.2. ATZ mitigates cardiac Aβ accumulation and vascular deposition in the Tg2576 heart

To evaluate the impact of ATZ on cardiac amyloid-β pathology, we analyzed the levels and spatial distribution of Aβ in the heart of WT, Tg2576, and ATZ-treated Tg2576 mice (Fig. 2). Immunofluorescence staining revealed a significant accumulation of Aβ in the hearts of Tg2576 mice compared to WT controls, which was remarkably attenuated upon chronic ATZ administration (Fig. 2A). Dual-labeling spatial analysis further localized these Aβ aggregates directly to the cardiac vasculature (Fig 2A). Specifically, Tg2576 mice exhibited severe concentric Aβ deposition spanning the endothelial lining (CD31+) and the surrounding vascular smooth muscle layer (α-SMA+) of cardiac vessels, suggestive of cardiac vascular amyloid pathology. Importantly, ATZ treatment robustly reduced this vascular and intramural Aβ burden. Image quantification confirmed a significant increase in the percentage of overall cardiac Aβ area in Tg2576 mice, which was profoundly rescued by ATZ therapy (Fig. 2C). To validate these histological findings at the biochemical level, we assessed the deposition of Aβ species by WB within whole heart lysates. Immunoblot analysis identified a sizeable increase in Aβ dimers, a critical driver of localized cytotoxicity, in Tg2576 hearts (Fig. 2B-D). Additionally, ELISA assays revealed significant elevations in cardiac Aβ40 levels (the most vasculotropic Aβ peptide) within both untreated and ATZ-treated Tg2576 cohorts relative to WT mice, while cardiac Aβ42 trends remained statistically non-significant across groups (Fig. 2E-F).

**Figure 2.**
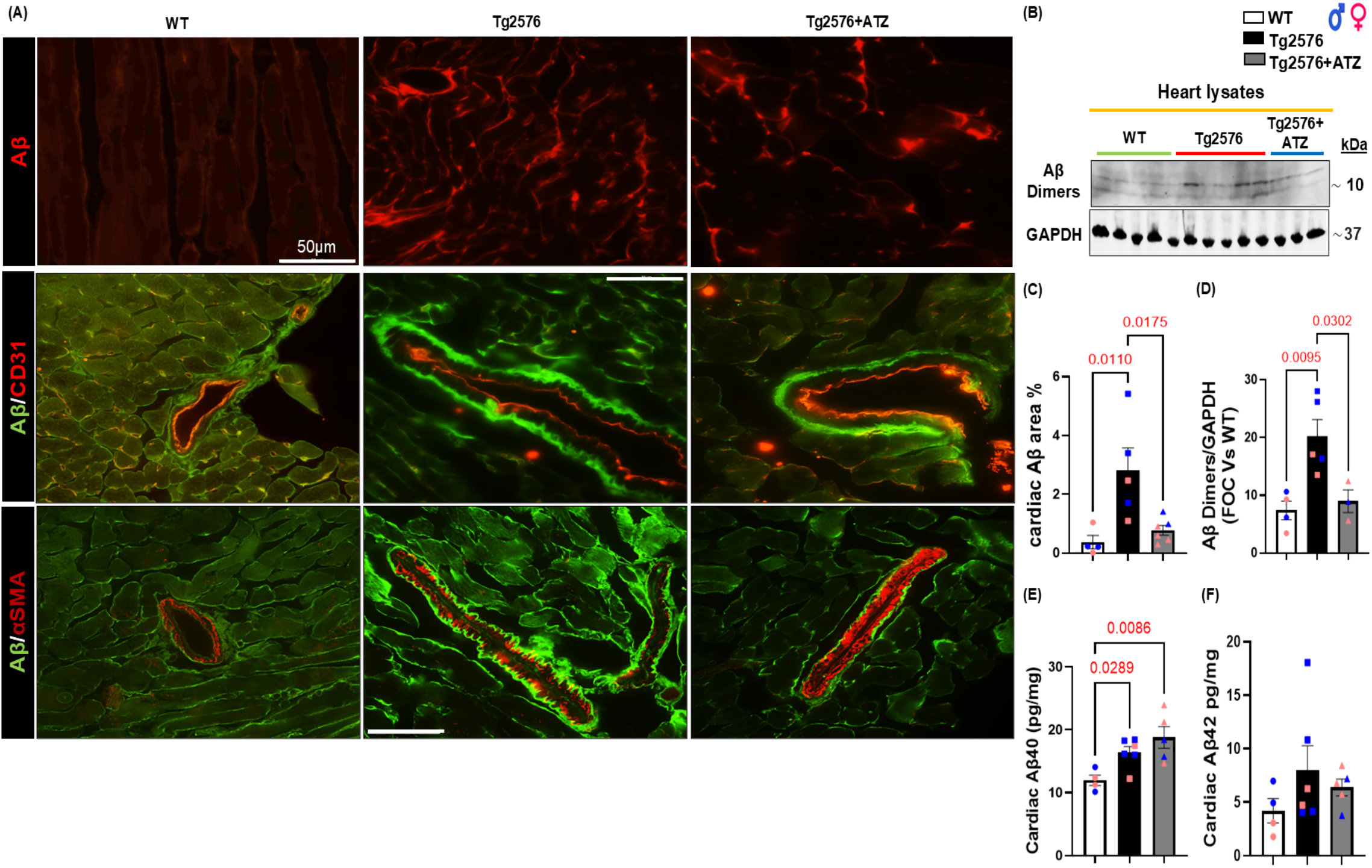
Acetazolamide (ATZ) reduces cardiovascular Aβ accumulation in Tg2576 mice. (**A**) Representative immunofluorescence images of cardiac tissue sections from WT, Tg2576, and Tg2576+ATZ mice. Top row: Aβ staining (red) showing parenchymal deposition (scale bar: 50 µm). Middle row: Co-staining of Aβ (green) and the endothelial marker CD31 (red). Bottom row: Co-staining of Aβ (green) and the vascular smooth muscle marker α-smooth muscle actin (αSMA, red; scale bar: 50 µm). (**B**) Representative Western blot analysis of Aβ dimers in heart lysates across the experimental groups, with GAPDH serving as the loading control. (**C**) Quantitative analysis of IHC cardiac Aβ area percentage (WT n=4; Tg2576 n=5; Tg2576+ATZ n=6). (**D**) Densitometric quantification of cardiac Aβ dimers normalized to GAPDH, expressed as (WT n=4; Tg2576 n=5; Tg2576+ATZ n=3). (**E**-**F**) ELISA quantification of cardiac Aβ40 (**E**) and cardiac Aβ42. (**F**) Concentrations expressed in pg/mg of total protein (WT n=4; Tg2576 n=6; Tg2576+ATZ n=5). Individual data points are color-coded by sex (blue = males; pink = females). Data are presented as mean±SEM. Statistical significance was determined using One-way ANOVA with Tukey’s post hoc test; exact p-values are indicated above the histograms.

Overall, these data provide evidence that ATZ treatment mitigates cardiac Aβ dimers and vascular Aβ accumulation within the myocardial parenchyma of a transgenic mouse model of cerebral amyloidosis and AD.

### 2.3. ATZ treatment preserves cardiac neurotrophic signaling and myocardial innervation in Tg2576 mice by rescuing the BDNF/TrkB/CREB axis and p75NTR maladaptive activation

AD-Aβ pathology profoundly disrupts neurotrophic signaling by depleting BDNF (Aggarwal, Kondeti et al. 2013, Chen, Sawa et al. 2018, Numakawa and Odaka 2021, Gao, Zhang et al. 2022, Xue, Waseem et al. 2022, Shen, Ardianto et al. 2023, Elia, Parodi-Rullan et al. 2026), a critical regulator of neuronal survival, synaptic function, and cardiac homeostasis (Aggarwal, Kondeti et al. 2013, Pius-Sadowska and Machalinski 2017, Miranda, Morici et al. 2019, Di Rosa, Zimbone et al. 2021, Elia, Cannavo et al. 2021, Hang, Zhu et al. 2021, Maroofi, Bagheri Rouch et al. 2022). Recent work has shown that Aβ-driven deterioration of BDNF signaling extends beyond the CNS, impairing cardiac innervation and myocardial neurotrophic support, thereby exacerbating cardiac dysfunction (Elia and Fossati 2023, Elia, Parodi-Rullan et al. 2026). We have demonstrated that the FDA-approved CAIs ATZ and MTZ exert potent neurovascular protective effects in AD/CAA models by preventing Aβ-induced mitochondrial apoptosis, reducing cerebrovascular amyloid deposition, attenuating neuroinflammation, and preserving cognitive function (Fossati, Giannoni et al. 2016, Solesio, Peixoto et al. 2018, Lemon, Canepa et al. 2021, Anzovino, Canepa et al. 2023, Canepa, Parodi-Rullan et al. 2023, Gutierrez-Jimenez, Rasmussen et al. 2025). Despite their established efficacy and safety in other cardiovascular diseases, such as hypertension (Hoff, Zou et al. 2020, Garcia-Llorca, Carta et al. 2024), or heart failure (Mullens, Dauw et al. 2022, Augusto, Martens et al. 2025), their impact on cardiac neurotrophic signaling and innervation in AD-associated heart pathology remains unknown. To address this gap, we explored whether ATZ treatment could mitigate AD-mediated impairments in cardiac neuro-signaling. Immunoblot evidence showed a significant reduction in the expression of BDNF, neuronal regeneration marker GAP-43, and sympathetic marker dopamine β-hydroxylase (DβH) in the Tg2576 hearts compared with age-matched WT mice (Fig. 3A-B), while long-term ATZ treatment remarkably rescued the levels of all these neuromodulators, suggesting significant preservation of cardiac neurotrophic support and heart neuronal integrity. Histological examinations further validated the biochemical results, demonstrating a reduction in tyrosine hydroxylase-positive (TH+) sympathetic fibers and GAP-43+ regenerating nerve endings within the Tg2576 myocardium. Critically, ATZ administration significantly recovered both TH+ and GAP-43+ fiber density, revealing effective restoration of cardiac innervation and neuronal remodeling (Fig. 3C-E).

**Figure 3.**
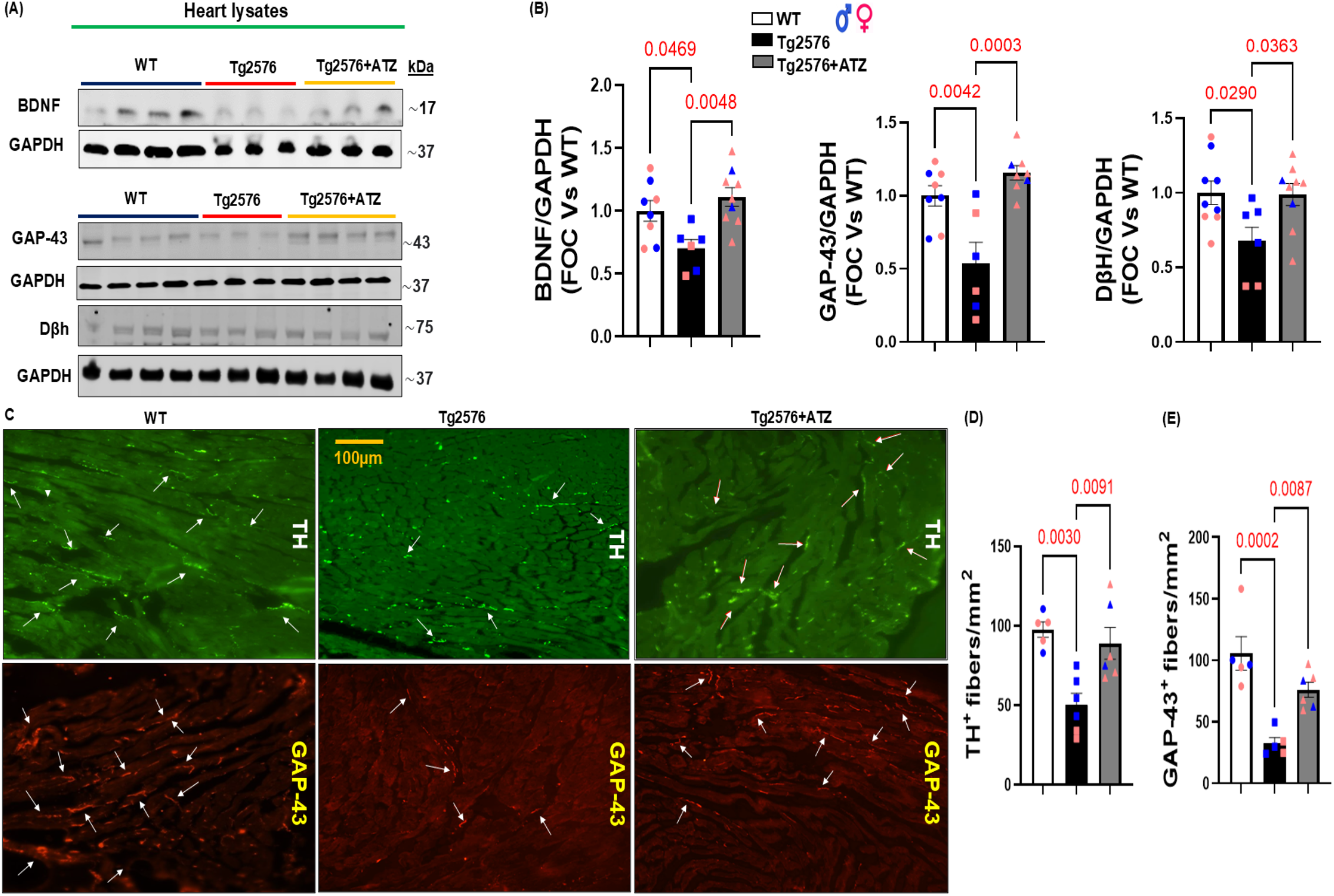
ATZ rescues Aβ-induced dysregulation of cardiac neuro-signaling and myocardial innervation in the Tg2576-AD model. (**A**) Representative immunoblots of BDNF, GAP-43, and dopamine β-hydroxylase (DβH) in heart lysates from WT, Tg2576, and ATZ-treated Tg2576 mice. GAPDH served as the loading control. (**B**) Quantitative densitometric analyses demonstrating significant reductions in BDNF, GAP-43, and DβH expression in Tg2576 hearts compared with WT controls, all of which were significantly and fully restored by chronic ATZ treatment (WT n=9; Tg2576 n=7; Tg2576+ATZ n=9). (**C**) Representative immunofluorescence images showing myocardial tyrosine hydroxylase (TH)-positive (TH+) sympathetic nerve fibers (green) and GAP-43-positive regenerating fibers (red) in WT, Tg2576, and Tg2576+ATZ mice. Arrows indicate nerve fibers (scale bar: 100 µm). (**D**-**E**) Quantification of TH+ fiber density (**D**) and GAP-43+ regenerating fibers (**E**) demonstrating loss of cardiac innervation in Tg2576 mice and significant recovery following ATZ administration (WT n=5; Tg2576 n=6; Tg2576+ATZ n=6). Data are presented as mean±SEM. Individual data points represent independent biological replicates, color-and shape-coded by sex (blue = males; pink = females). Statistical significance was determined using a multi-group One-way ANOVA with Tukey’s post hoc test and is indicated in the panels.

To further characterize the molecular mechanisms underlying this neuroprotective response, we evaluated crucial mediators of the cardiac neurotrophic signaling cascade. Tg2576 hearts exhibited a clear decrease in mature BDNF and NGF levels, accompanied by a considerable increase in proBDNF (Supplemental Fig. 6), suggesting a pathological neurotrophic signaling imbalance, consistent with that found in the brain in AD and other neurodegenerative diseases (Fahnestock, Michalski et al. 2001, Pedraza, Podlesniy et al. 2005). Remarkably, chronic ATZ therapy significantly rescued cardiac mature neurotrophin levels while reducing proBDNF. Aligning with cardiac mature BDNF reduction, Tg2576 hearts displayed considerably reduced phosphorylated TrkB (the active BDNF receptor form) and decreased expression of both total and phosphorylated CREB levels, the major downstream transcription factor of BDNF signaling (Fig. 4A-B). ATZ significantly rescued TrkB activation and CREB signaling, suggesting re-establishment of the cardioprotective BDNF/TrkB/CREB axis. Simultaneously, the untreated Tg2576 group showed a marked elevation in p75NTR expression, a receptor highly implicated in degenerative and apoptotic responses mediated by Aβ in AD pathology (Fig 4A, bottom), (Perini, Della-Bianca et al. 2002, Bruno, Abondio et al. 2023, Shanks, Chen et al. 2024), while this p75NTR upregulation was not present in Tg2576 mice treated with ATZ. Consistent with the in vivo findings, oligomeric Aβ40 exposure significantly increased p75NTR expression in AC16 cardiomyocytes, whereas ATZ treatment significantly attenuated this response (Fig. 4C). AC16 cells co-treated with an analog CAI, MTZ, also showed some reduction in p75NTR, although not significant. These findings support the involvement of p75NTR in Aβ-induced cardiac cellular stress and pro-apoptotic signaling, extending previous evidence of direct Aβ–p75NTR interactions largely established in neuronal cells (Sotthibundhu, Sykes et al. 2008, Knowles, Simmons et al. 2013, Demuth, Hosseini et al. 2023), and highlighting for the first time the positive effects of CAIs against p75 maladaptive activation. These *in vitro* observations were further corroborated in Tg2576 hearts by histological analyses, which revealed substantial p75NTR upregulation within myocardial and vascular compartments, with prominent colocalization with areas of Aβ accumulation (Fig. 4D-E). Notably, these IHC experiments confirmed that ATZ significantly reduced p75NTR expression and decreased its vascular deposition.

**Figure 4.**
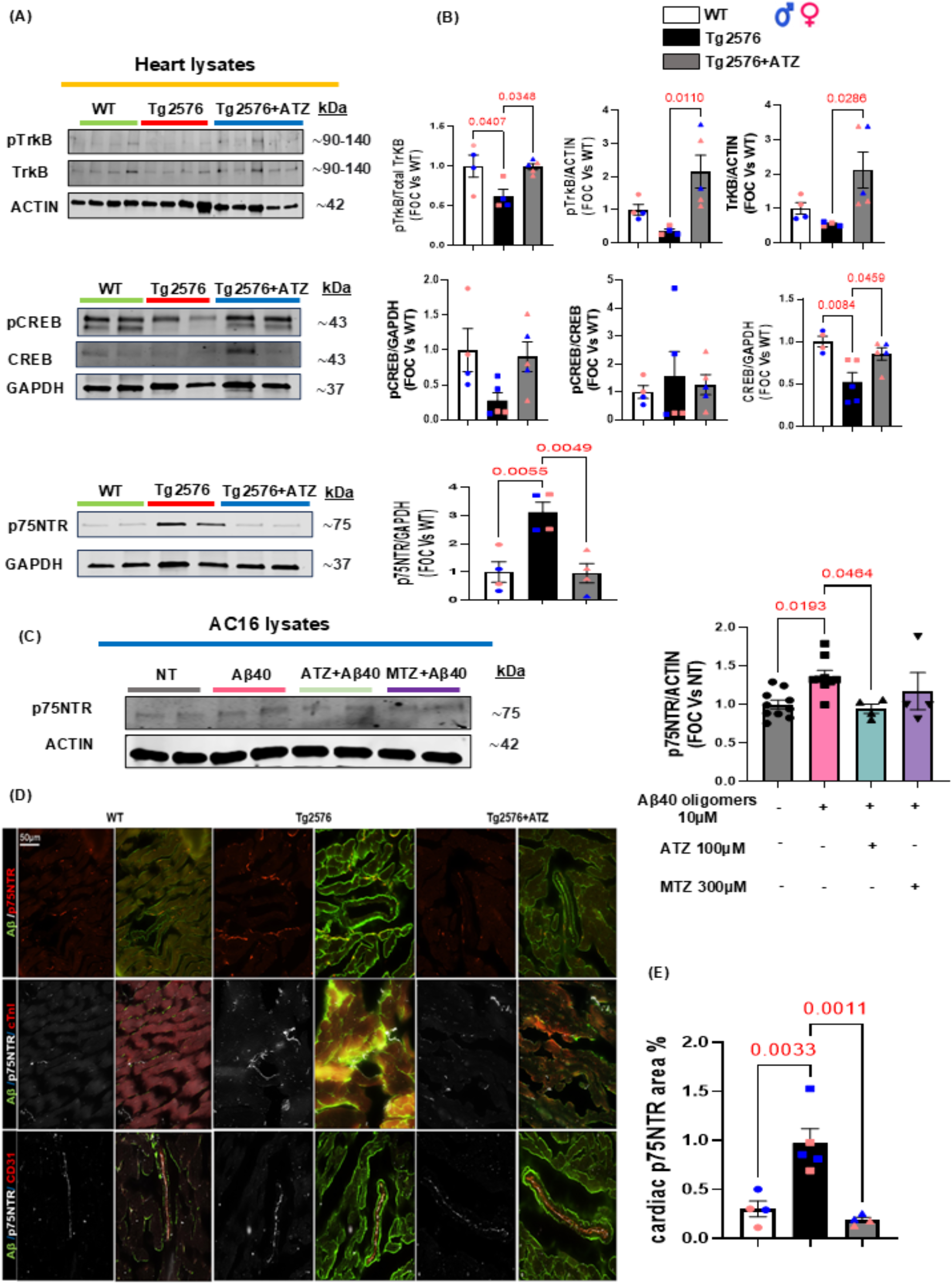
Acetazolamide restores cardiac BDNF–TrkB–CREB signaling and suppresses p75NTR activation in Tg2576 hearts. (**A**) Representative immunoblots of phosphorylated TrkB (pTrkB), total TrkB, phosphorylated CREB (pCREB), total CREB, and p75NTR in cardiac lysates from WT, Tg2576, and Tg2576+ATZ mice. (**B**) Densitometric analyses demonstrating significant reductions in TrkB activation, CREB phosphorylation, and total CREB expression in Tg2576 hearts, accompanied by robust p75NTR upregulation (WT n=4; Tg2576 n=5; Tg2576+ATZ n=5). (**C**) Representative immunoblots (upper panels) and densitometric quantitative analysis (lower panel) showing protein levels of p75NTR in total protein lysates from human cardiomyocytes (AC16) stimulated with human Aβ40 oligomers (10 μM) for 16 h. Before Aβ40 oligomer treatment, cells were pretreated with either ATZ (100 μM) or MTZ (300 μM) for 1 h. ACTIN levels were used as a loading control. (n = 4-10). (NT = Not Treated). (**D**) Representative digital images showing colocalization of p75NTR (red) with amyloid-β deposits (green) within myocardial tissue and vascular structures. Additional staining with cardiac troponin I (cTnI) and CD31 identifies cardiomyocyte and endothelial compartments, respectively. Tg2576 hearts exhibit robust p75NTR accumulation associated with amyloid buildup in myocardial and vascular regions, whereas ATZ treatment markedly attenuates p75NTR expression (scale bar: 50 µm). (**E**) Quantitative analysis of cardiac p75NTR-positive area demonstrating significant elevation in Tg2576 mice and substantial suppression following chronic ATZ treatment. Data are presented as mean±SEM (WT n=4; Tg2576 n=5; Tg2576+ATZ n=4). Statistical significance is indicated in the panels. Chronic ATZ administration restored TrkB and CREB signaling while markedly suppressing p75NTR expression. Data are presented as mean±SEM. Statistical significance was determined using a One-way ANOVA with Tukey’s post hoc test, and exact p-values are indicated in the panels.

Collectively, these findings confirm a robust cardiac neurotrophic deterioration as a distinct feature of AD-related myocardial pathogenesis and establish that ATZ rescues cardiac neurotrophin homeostasis, maintaining myocardial innervation, re-establishing the BDNF/TrkB/CREB signaling axis, and limiting maladaptive p75NTR overexpression, which is known to be linked to its activation (Bhakar, Howell et al. 2003).

### 2.4. Carbonic anhydrase inhibitors preserve the BDNF/TrkB/CREB neurotrophic axis in AC16 Cardiomyocytes and Neuroblastoma Cells

Our *in vivo* results suggest that ATZ treatment prevents cardiac neuro-signaling dysregulation and preserves myocardial innervation against the detrimental effects of Aβ in Tg2576 mice. Thus, to confirm the therapeutic potential of CAIs against Aβ-mediated neurodegeneration, we explored their capacity to rescue neurotrophic dysfunction and neuronal sprouting in SH-SY5Y human neural cells exposed to toxic Aβ40 and Aβ42 oligomers (Fig. 5 and Supplemental Fig. 1). Both Aβ40 and Aβ42 significantly affected neurotrophic modulators in SH-SY5Y cells, as evidenced by a significant decrease in the protein expression levels of BDNF and GAP-43, compared to non-treated (NT) controls (Fig. 5A-B). Interestingly, ATZ exposure completely rescued Aβ-induced neurotoxicity, restoring both GAP-43 and BDNF expression. We did not observe significant changes upon MTZ treatment, thus suggesting a preferential pharmacological efficacy of ATZ in preserving neurotrophic support in neuronal cells. In line with our previous molecular findings (Elia, Parodi-Rullan et al. 2026), Aβ40 and Aβ42 oligomers also triggered severe neurite dystrophy, resulting in a significant reduction in total neurite length (Supplemental Fig. 1A-D), which was significantly restored by ATZ treatment.

**Figure 5.**
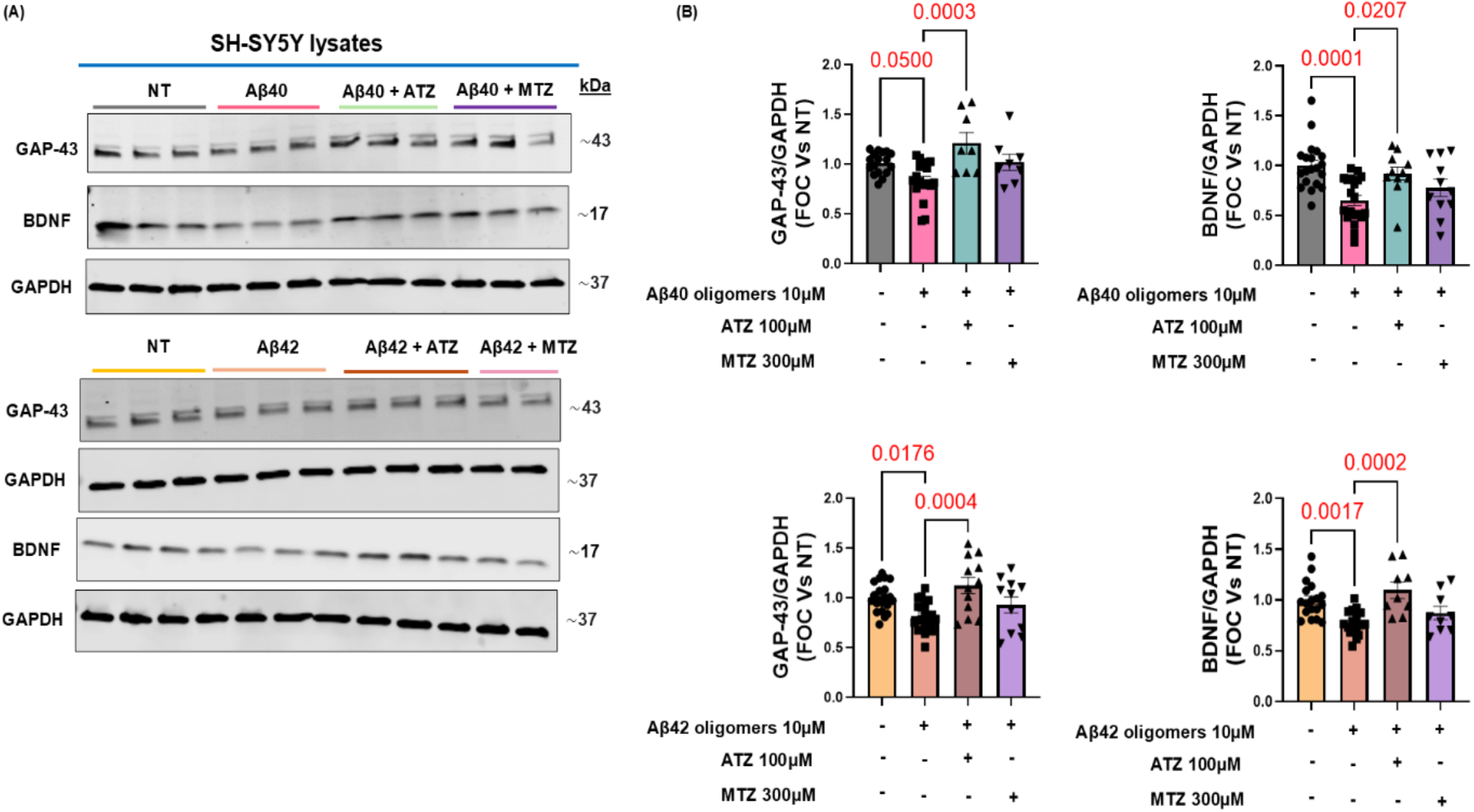
ATZ treatment prevents GAP-43 and BDNF expression in SH-SY5Y cells exposed to Aβ oligomers. (**A**) Representative Western blot images showing GAP-43 and BDNF protein expression in SH-SY5Y cell lysates following treatment with Aβ40 or Aβ42 oligomers (10 μM) for 16 h. Before Aβ40 or Aβ42 oligomer treatment, cells were pretreated with either ATZ (100 μM) or MTZ (300 μM) for 1 h. (**B**) Densitometric quantification of GAP-43 and BDNF protein levels normalized to GAPDH and expressed as Fold of Change (FOC) relative to the Not Treated (NT) control group. (n=8-21). Data are presented as mean±SEM of independent experiments (represented by individual data points). Exact P-values are denoted above the brackets indicating statistical comparisons determined by One-way ANOVA followed by post hoc analysis. (NT = Not Treated).

To determine whether CAIs also directly protect cardiomyocytes from Aβ-driven neurotrophic impairment, human AC16 cardiomyocytes were challenged with Aβ40 oligomers in the presence or absence of ATZ or MTZ. Immunoblot analyses confirmed that Aβ40 exposure significantly decreased BDNF expression and induced a marked increase in cleaved caspase-3 levels, supporting an activation of the apoptotic signaling pathway in cardiomyocytes (Fig. 6A-B) (Elia, Parodi-Rullan et al. 2026). In these cells, both ATZ and MTZ mitigated Aβ-induced caspase-3 activation and MTZ treatment significantly preserved BDNF expression, while the effects of ATZ on BDNF were not significant, suggesting a possible cell-and pathway-selective action of different CAIs.

**Figure 6.**
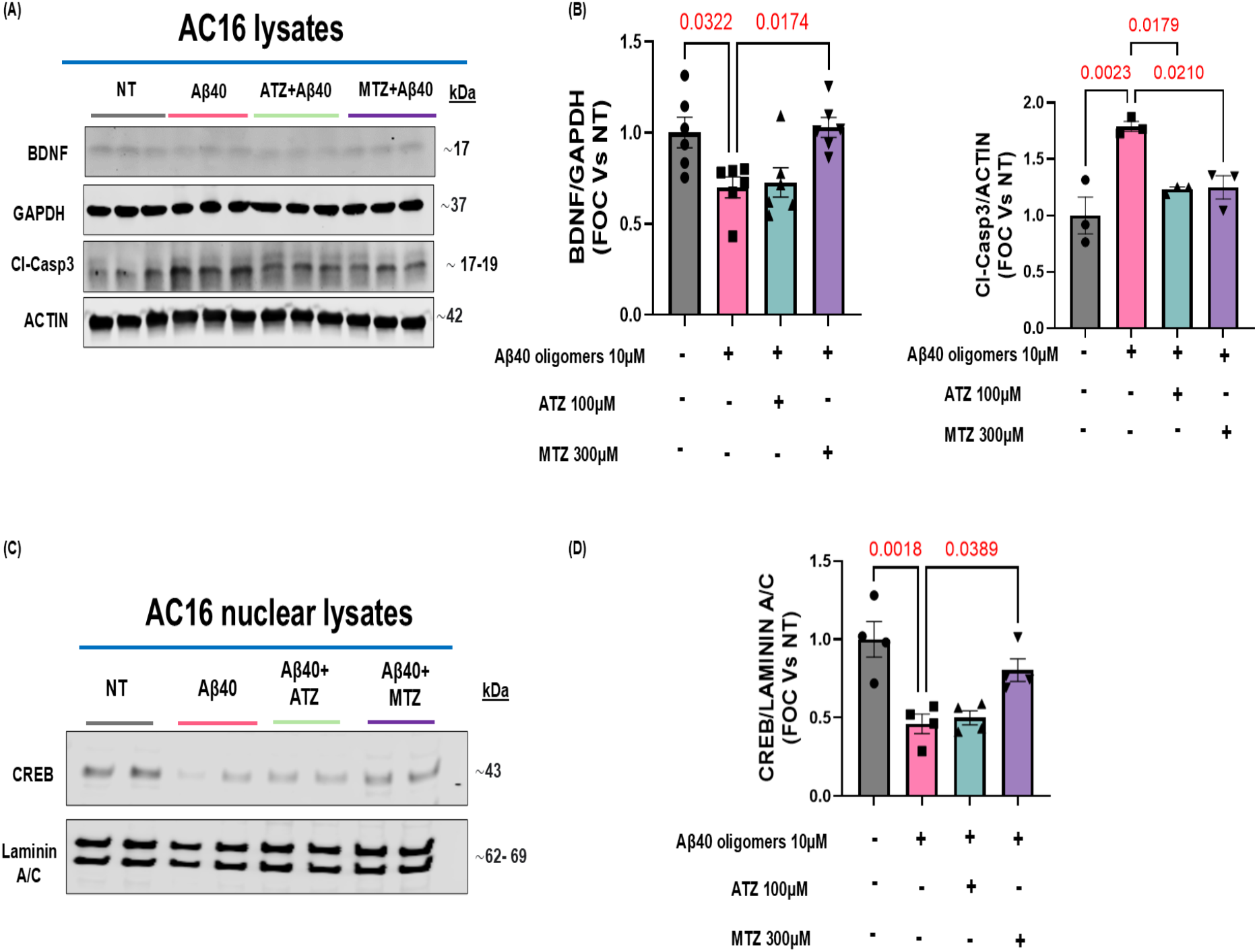
Carbonic anhydrase inhibitors preserve BDNF–CREB signaling and attenuate apoptotic activation in Aβ40-treated human AC16 cardiomyocytes. (A) Representative immunoblots of BDNF and cleaved caspase-3 (Cl-Casp3) in AC16 cell lysates following exposure to Aβ40 oligomers (10 μM) for 16 h. Before Aβ40 oligomer treatment, cells were pretreated with either ATZ (100 μM) or MTZ (300 μM) for 1 h. GAPDH and ACTIN were used as loading controls. (B) Quantification of BDNF and Cl-Casp3 protein levels normalized to respective loading controls. Aβ40 significantly reduced BDNF expression and increased apoptotic signaling, whereas CAIs treatment with MTZ restored neurotrophic support and both ATZ and MTZ attenuated caspase-3 activation (n=3-6). (C) Representative immunoblots of nuclear CREB in AC16 cardiomyocytes stimulated with Aβ40 oligomers (10 μM) for 15 min. Lamin A/C served as the nuclear loading control. Before Aβ40 oligomer treatment, cells were pretreated with either ATZ (100 μM), or MTZ (300 μM) for 10 min. (**D**) Quantitative analysis demonstrating significant preservation of CREB nuclear levels following MTZ treatment (n=4). Data are presented as mean±SEM. One-way ANOVA with Tukey’s multiple comparisons post hoc test: *P *<* 0.05 vs NT. (NT = Not Treated). Statistical significance is indicated in the figure.

To understand the molecular underpinnings behind Aβ-induced BDNF downregulation, we dissected the BDNF/TrkB/CREB signaling axis. Due to the dual role of CREB as a key downstream effector of BDNF-mediated survival signaling, and as a major transcription factor for BDNF, we evaluated acute changes in nuclear CREB levels in AC16 cardiomyocytes. Notably, Aβ40 oligomers induced a decline in nuclear CREB levels at 15 minutes, in line with acutely impaired neuro-signaling activity (Fig. 6C-D) and with our previous findings (Elia, Parodi-Rullan et al. 2026). Importantly, MTZ co-treatment robustly rescued nuclear CREB levels, in line with the rescue in BDNF expression. Complementary immunofluorescence investigations further confirmed this evidence (Supplementary Fig. 2). Aβ40-treated AC16 cells exhibited a substantial decrease in intracellular BDNF immunoreactivity, while CAIs, and more obviously MTZ, significantly preserved BDNF expression (Supplementary Fig. 2A), in line with the WB results. Likewise, Aβ40 exposure strikingly reduced nuclear CREB immunostaining, whereas CAIs intervention rescued CREB nuclear localization and expression (Supplementary Fig. 2B).

Altogether, these findings indicate that CAIs strongly prevent Aβ-induced cardiomyocyte impairment by preserving the BDNF/TrkB/CREB neurotrophic axis in both neurons and cardiomyocytes and counteracting apoptotic signaling, adding mechanistic support that carbonic anhydrase inhibition promotes efficient cardioprotective and neurotrophic effects against amyloid toxicity.

## 3. Discussion

The current investigation provides compelling evidence that carbonic anhydrase inhibition is a promising therapeutic approach to counteract cardiac AD-mediated maladaptive remodeling and neurotrophic factor impairment. Although the harmful impact of Aβ accumulation on the central nervous system has been widely described, its effect on the cardiac neuro-signaling network had remained uninvestigated until recently. Our recent study (Elia, Parodi-Rullan et al. 2026) showed that progressive myocardial Aβ buildup in aged Tg2576 mice induces extensive cardiac dysfunction characterized by myocardial fibrosis, cardiomyocyte hypertrophy, ventricular impairment, vascular amyloid-related inflammation, caspase activation, and significant cardiac neurotrophic signaling dysregulation. It has been demonstrated that CAIs promote cerebrovascular homeostasis, revert the neuroinflammatory state, favor Aβ and tau clearance, and hamper cognitive impairment in models of cerebral amyloidosis and AD (Fossati, Giannoni et al. 2016, Solesio, Peixoto et al. 2018, Canepa, Parodi-Rullan et al. 2023, Carlsen, Fossati et al. 2025, Gutierrez-Jimenez, Rasmussen et al. 2025, Lopez, Siddiqi et al. 2025). Importantly, a very recent study, using machine learning approaches, has shown that ATZ, the CAI used in our *in vivo* study, is among the 3 most promising FDA-approved drugs that could reverse gene expression perturbations induced by AD pathology and, strikingly, its use was associated with significantly reduced risk for AD in human subjects (Vitali, Raikes et al. 2026).

Here, we hypothesized that CAIs’ treatment may ameliorate cardiac amyloid pathology and heart function, by both enhancing cerebrovascular amyloid clearance, as we have previously shown (Canepa, Parodi-Rullan et al. 2023), therefore reducing amyloid aggregates released into the periphery and accumulating in the heart, as well as by directly reducing caspase activation, as previously shown by our lab in both cerebral vascular and neural cells (Fossati, Giannoni et al. 2016, Solesio, Peixoto et al. 2018, Anzovino, Canepa et al. 2023), thus ameliorating cardiomyocyte and cardiac neuronal health. Consistent with this hypothesis, the present study shows that chronic intervention with the FDA-approved carbonic anhydrase inhibitor ATZ profoundly decreased myocardial Aβ accumulations, alleviated vascular and interstitial amyloid-associated pathology, repressed caspase 3 activation in cardiomyocytes and SAA3 cardiovascular infiltration, rescued heart neurotrophic fitness and myocardial innervation and, overall, significantly enhanced cardiac function in Tg2576 mice. This *in vivo* evidence was further corroborated by complementary human neural and cardiac cellular models, providing mechanistic support that carbonic anhydrase inhibition effectively prevents Aβ-mediated neurotrophic dysfunction.

While confirming prior results (Elia, Parodi-Rullan et al. 2026), this study added further mechanistic findings to demonstrate that Aβ deposition deranges the myocardial BDNF/TrkB/CREB axis, promoting a pathological shift toward pro-neurotrophin/p75NTR pro-apoptotic signaling, and pronounced adrenergic denervation and regenerative nerve fibers depauperization, ultimately impairing cardiac function. Notably, CA inhibition rescued all these pathways both *in vitro* and *in vivo*.

ATZ robustly limited Aβ-induced neurotoxicity alongside neuronal sprouting remodeling, preserving expression of the neuronal regeneration marker GAP-43 and the essential neurotrophin BDNF in SH-SY5Y neural cells. Aβ40 oligomers challenge of human AC16 cardiomyocytes culminated in a robust downregulation of CREB activation and BDNF expression, as well as an increase in maladaptive p75NTR expression, and these effects were counteracted by the CAIs ATZ and/or MTZ. Interestingly, ATZ appeared to have better effects in restoring BDNF in neural cells, while MTZ seemed to prevent BDNF loss more efficiently in cardiomyocytes, pointing to cell-selective neurotrophic actions of these compounds. However, both ATZ and MTZ significantly reduced Aβ-induced caspase activation in cardiomyocytes, validating the anti-apoptotic effects that we have previously shown in neural and cerebrovascular cells (Solesio, Peixoto et al. 2018, Anzovino, Canepa et al. 2023).

Altogether, these data establish the cardiac BDNF/TrkB/CREB and p75 axes as key downstream targets of Aβ toxicity and place carbonic anhydrase inhibition as a compelling cardio-neuroprotective therapy to halt the vicious cycle linking amyloid pathology, neuro-signaling deregulation, apoptosis, and cardiac deterioration. We also provide evidence for a potential positive feedback loop in the AD cardiac tissue: Aβ binds/activates p75NTR, and this activation in turn can drive further p75NTR (and co-receptor) expression and caspase activation, consistent with the dependence receptor model where p75NTR itself becomes a hub for amyloid toxicity in AD (Chakravarthy, Gaudet et al. 2010, Saadipour, Yang et al. 2013). Additionally, this study aligns with previous literature, showing that pathological Aβ levels acutely impair functional CREB activity. This effect was previously shown to be occurring due to blocking of the PKA-CREB pathway or trapping of phosphorylated CREB outside the nucleus (Vitolo, Sant’Angelo et al. 2002, Arvanitis, Ducatenzeiler et al. 2007), which is also consistent with our data showing lower amounts of nuclear CREB in Aβ-challenged cardiomyocytes acutely, as early as 15 minutes after challenge.

Our results also show that ATZ treatment reduces deposits of SAA3, a recognized marker of cardiovascular inflammation in ischemic heart disorders (Thompson, Wilson et al. 2018, Chait, den Hartigh et al. 2020, den Hartigh, May et al. 2023, Cimini, Hansmann et al. 2025), within the vascular walls of Tg2576 hearts. It is well-established that Aβ oligomers are able to promote the aggregation of heterologous amyloid species through amyloid seeding mechanisms (Ulm, Borchelt et al. 2021), escalating cellular stress, inflammation, and tissue remodeling across multiple cell types (Parodi-Rullan, Javadov et al. 2021, Jang, Chapa-Dubocq et al. 2022). Hence, the reduction of cardiac SAA3, as well as cardiac Aβ accumulation, could be linked to a reduction of CAA pathology induced by CAIs, which would also reduce BBB leakage and therefore the amount of Aβ oligomers able to escape the brain towards the peripheral circulation and eventually the heart. Nevertheless, the role of Aβ oligomers in fostering systemic and cardiac seeding of other amyloids remains speculative and is not directly demonstrated in the current report.

Remarkably, in previous studies, CAIs have exhibited efficacy in reducing cerebral amyloid pathology and CAA, cerebrovascular/cognitive impairment, and neuroinflammation in AD, CAA or tauopathy models (Fossati, Giannoni et al. 2016, Solesio, Peixoto et al. 2018, Lemon, Canepa et al. 2021, Anzovino, Canepa et al. 2023, Canepa, Parodi-Rullan et al. 2023, Gutierrez-Jimenez, Rasmussen et al. 2025), and an exciting recent report employing machine learning approaches has identified ATZ as one of the three most promising drugs to repurpose for AD therapy (Vitali, Raikes et al. 2026) Aligning with these studies, our current findings significantly expand CAIs’ therapeutic portfolio and support a distinct dual-action strategy that simultaneously targets both central and cardiovascular manifestations of AD pathology. The translational relevance of these observations is further strengthened by the fact that ATZ and MTZ are FDA-approved medications with a well-established safety regimen and known BBB permeability (Provensi, Carta et al. 2019, Lemon, Canepa et al. 2021). This supports the potential to repurpose CAIs as a new therapeutic strategy not only to reduce AD cerebral pathology and detrimental cognitive effects, but also to ameliorate AD-related cardiac changes, emphasizing the importance of additional studies in this direction.

While this contribution offers meaningful mechanistic and translational insights into AD cardiac effects and the potential protective role of CAIs, several limitations should be taken into consideration.

Invasive cardiac pressure-volume loop analysis was not performed because aged Tg2576 mice exhibit increased frailty and altered physiological responses to anesthesia, which can introduce significant cardiovascular and autonomic confounders. Recent studies have demonstrated genotype-specific instability during anesthesia and evidence of autonomic dysfunction in AD mouse models, supporting the use of less invasive approaches when possible. To minimize procedural stress and preserve animal welfare in this vulnerable aging cohort, cardiac function was assessed using echocardiography and complementary molecular analyses. While such examinations could have supplied further information related to cardiac performance, they also would have considerably increased the risk of severely impacting animal lifespan and experimental integrity (Miranda-Silva, Sequeira et al. 2021, Chen, Kwok et al. 2023, Bratke, Schmid et al. 2024).

Although our findings suggest BDNF/TrkB/CREB axis derangement as a key feature of AD-associated cardiac pathology, other NTFs, including NT-3 and NT-4 signaling, may also participate in myocardial innervation, neurovascular health, and cardiac function (Donovan, Hahn et al. 1996, Story, Dicarlo et al. 2000, Aggarwal, Kondeti et al. 2013, Patsy, Ge et al. 2025). Future studies analyzing the interconnections among multiple neurotrophic systems will be valuable for defining a more extensive scaffold of neuro-signaling dysregulation within the AD myocardium.

While BDNF/TrkB/CREB and p75NTR signaling rescue was robustly associated with CAIs-driven cardioprotective effects, conclusive mechanistic confirmation will require targeted loss-of-function strategies entailing BDNF, TrkB, and p75NTR to fully establish causal pathway dependency. However, the accomplishment of genetic interventions, including cardiomyocyte-specific BDNF deletion in aged Tg2576 mice, poses notable challenges, such as age-dependent toxic effects and the need for comprehensive longitudinal monitoring (Jankowsky and Zheng 2017, Zhong, Peng et al. 2024). Based on the robust recovery of cardiac BDNF signaling observed in the present study, we also hypothesize that direct TrkB activation may represent a viable therapeutic strategy to counteract AD-associated cardiac maladaptive remodeling and functional decline. Yet, rigorous *in vivo* validation of this mechanism remains warranted and constitutes a direction for future investigation.

Further validation across independent AD models would support the generalizability and translational relevance of our observations. Although Tg2576 mice are a well-characterized model of cerebral amyloidosis with CAA (Hsiao, Chapman et al. 1996), they do not completely mimic the multifactorial nature of the human AD phenotype or the complexity of age-associated cardiovascular disease. Expanding these findings to complementary models, such asother models of CAA, ,3xTg-AD mice or tauopathy models, will therefore be crucial to characterize the more comprehensive applicability and strength of the observed cardiac phenotype. Additionally, our previous studies have confirmed the dysregulation of these cardiac pathways in human AD post-mortem heart tissue (Elia, Parodi-Rullan et al. 2026). However, the potential protective effects of ATZ or MTZ on the human heart in MCI or AD patients can only be assessed through dedicated clinical trials, which we expected to initiate in the close future.

Crucially, to our knowledge, the present research provides the first evidence and characterization of the positive effects of FDA-approved CAIs against cardiac abnormalities in an AD model, supporting a novel beneficial pharmacological role of CAI medications to prevent AD-mediated cardiac neuro-signaling remodeling, myocardial denervation, and heart dysfunction. Extending these observations across further AD models and different cardiac pathologies remain a priority, but demands prolonged aging, extensive colony maintenance, and considerable logistical assets.

Ultimately, several therapeutic questions remain, including optimal dosing protocols, treatment timing, long-term effectiveness, robustness of response, and potential sex-dependent effects.

Despite these limitations, the present investigation provides a solid foundation for the novel beneficial role of FDA-approved CAIs, such as ATZ, to rescue AD-mediated cardiac pathology, and highlights the BDNF/TrkB/CREB and p75NTRsignaling axes as promising therapeutic targets to prevent or correct both central and cardiovascular complications induced by AD and amyloidosis, while emphasizing the need for additional mechanistic, longitudinal, and cross-model analyses.

## 4. Conclusions

In conclusion, our work provides the first exhaustive evidence that carbonic anhydrase inhibition ameliorates AD-related cardiac neuro-signaling and myocardial nervous system impairments. We validate the BDNF/TrkB/CREB signaling axis as a central molecular target of Aβ-mediated cardiac remodeling and display that long-term carbonic anhydrase inhibition valuably preserves this pathway, represses maladaptive p75NTR signaling, prevents myocardial denervation, and decreases cardiac amyloid load, thus enhancing cardiac function. These findings demonstrate a previously unknown cardio-neuroprotective role of CAIs and ascertain the heart BDNF/TrkB/p75 axis as a novel therapeutic target in AD-associated cardiovascular disease. With their documented clinical benefit and promising safety profile, FDA-approved CAIs such as ATZ can be positioned as highly unique candidates for drug repurposing strategies aimed at simultaneously addressing neurodegenerative, cerebrovascular, and cardiac manifestations of AD and amyloid diseases. Finally, our data serve as a solid rational ground to support clinical trials aimed at considering CAIs as a novel therapeutic target against AD-related cardiomyopathy and Aβ-induced cardiac neurotrophic system remodeling, which could potentially improve patient outcomes and quality of life while easing the healthcare system’s burden.

## 5. Experimental Section

### 5.1 Animal Model and Therapeutic Intervention

Male and female Tg2576 mice, a model of cerebral amyloidosis bearing the Swedish mutation (KM670/671NL), and age-matched wild-type (WT) littermates were bred and genotyped in-house. Mice were maintained under controlled conditions (∼22°C and in a 12-hour light-dark cycle, lights from 7 am to 7 pm) with unrestricted access to food and water. The generation of B6; SJL-Tg (APPSWE)2576Kha mice (Tg2576) on a B6; SJL Mixed Background was described in (Hsiao, Chapman et al. 1996) . Tg2576 mice are known to develop numerous cerebral parenchymal amyloid-β plaques at 11-13 months and CAA at 10-11 months (Robbins, Betensky et al. 2006). To assess the translational potential of CAIs in counteracting AD-driven cardiac pathology and neuro-signaling impairments, Tg2576 mice were administered an acetazolamide (ATZ)-enriched diet (20 mg/kg/day, ∼100 ppm; 5053 by TestDiets, Quakertown, PA) continuously from 6 to 13 months of age, using a regimen consistent with prior preclinical studies (Canepa, Parodi-Rullan et al. 2023, Gutierrez-Jimenez, Rasmussen et al. 2025). Body weight was measured biweekly during the first two months and again before cardiovascular assessments, while skin and fur health were monitored weekly to detect any signs of toxicity. This rigorous long-term dosing strategy confirms the safety of CAI treatment in this AD mouse model. All experiments and animal protocols were performed in equal numbers of male and female mice, according to procedures approved by the Institutional Animal Care and Use Committee of Temple University School of Medicine and conformed to the National Research Council Guide for the Care and Use of Laboratory Animals published by the US National Institutes of Health (2011, eighth edition).

### 5.2 *In vivo* transthoracic echocardiography of left ventricular function in murine models

Cardiac function was comprehensively assessed in all mouse cohorts, both prior to and following the onset of cerebral Aβ pathology via echocardiographic imaging. Specifically, transthoracic bi-dimensional M-mode echocardiography was blindly performed through the VisualSonics Vevo F2 (Visual Sonics) system (VisualSonics, Toronto, Canada) under controlled anesthesia, at baseline (4 months), one month after the beginning of the ATZ treatment at 8 months of age, and at the end of the study period (13 months old).

Before imaging, thoracic depilation was conducted with care to ensure optimal ultrasound contact. Mice were positioned in a clinostatic posture on a temperature-controlled platform equipped with embedded ECG electrodes, maintaining a stable body temperature of 37 °C. Anesthesia was delivered via an induction chamber using isoflurane (Zoetis IsoFlo, Kalamazoo, MI), with a 3.0% induction dose and a maintenance range of 1.0–2.5%, closely monitored to ensure physiological stability throughout the procedure. Cardiac structural and functional parameters, including LV wall thickness, end-diastolic diameter (LVEDD), end-systolic diameter (LVESD), end-diastolic volume (LVEDV), end-systolic volume (LVESV), ejection fraction (EF%), and fractional shortening (FS%), were quantified in accordance with established echocardiographic protocols (de Lucia, Wallner et al. 2019, Rigaud, Zarka et al. 2022). To detect subtle myocardial dysfunction, speckle tracking-based strain echocardiography (STE) was conducted using long-axis B-mode cine loops (300 frames at >200 fps), digitally recorded through VisualSonics VevoF2 system (VisualSonics, Toronto, Canada) and analyzed with Vevo LAB 1.7.1 software. LV endocardial deformation was quantified as peak strain (%) and strain rate (SR; 1/s) along both radial and longitudinal axes. Global and regional analyses were performed based on average measurements across six LV segments (basal, mid, apical anterior and posterior walls). Vector diagrams were generated to visualize the magnitude and direction of myocardial strain. Diastolic dysfunction was evaluated by calculating reverse global longitudinal strain (GLS) and global radial strain (GRS) rates, derived from long-axis B-mode traces as previously described (Schnelle, Catibog et al. 2018, de Lucia, Wallner et al. 2019). At the study endpoint (13 months), all animals were euthanized for organ harvesting and subsequent biomolecular and histological analyses.

### 5.3 Immunostaining and quantitative morphological assessment

Cardiac tissues were fixed in Zamboni’s solution (2% paraformaldehyde and picric acid) for 48 hours at 4°C, then sequentially cryoprotected in 10% sucrose in PBS for 1 hour and 20% sucrose in PBS for 24 hours at 4°C. Longitudinal sections (5 µm thick) were prepared using a Leica 2000R cryostat (Germany). Interstitial fibrosis was assessed with the Picrosirius Red Stain Kit (Abcam, ab150681), and stained sections were imaged using a Nikon NiE fluorescence microscope with a 20× objective (Cannavo, Marzano et al. 2019). Fibrotic area was quantified using automated threshold-based algorithms with ImageJ software (NIH, version 1.30) and expressed as the percentage of fibrotic tissue relative to the total myocardial area. All image analyses were performed by a single blinded operator to ensure consistency and minimize observer bias. To stain and quantify cardiac nerve fibers cardiac sections (5 µm thick) were processed using indirect immunofluorescence and associated proteins. Notably, adrenergic nerve fibers were labeled using rabbit polyclonal anti-tyrosine hydroxylase (TH; Millipore, AB152; 1:100), and regenerating nerve terminals were marked with anti-GAP-43 (Millipore, AB5220; 1:100). Sections were imaged using a Nikon Eclipse Ti fluorescence microscope with 20× and 60× objectives. Cardiac nerve endings appeared in a regular pattern along cardiomyocytes (de Lucia, Gambino et al. 2018). Nerve fiber density was quantified by overlaying a square grid (24 probes, 1 cm² each) on digital images using automated threshold-based algorithms through ImageJ software (NIH, version 1.30). Five randomly selected regions from three tissue sections per sample were analyzed. Intercepts between nerve fibers and the grid were counted, and nerve fiber density was expressed as mean intercepts per mm². All measurements were performed blindly by a single operator. To assess amyloid-β (Aβ) deposition, cardiac sections were immunostained with 6E10 anti-Aβ antibody (Covance, SIG-39320; 1:500) and co-stained with antibodies against cardiac troponin T (cTnT; Thermo Scientific, MA5-12960; 1:250), p75NTR (R&D Systems, AF1157; 1:100), cardiac troponin I (cTnI; Abcam, ab47003; 1:200), or cleaved caspase-3 (Cl-Casp-3; Cell Signaling, #94530; 1:200). Nuclei were counterstained with DAPI (Sigma-Aldrich, D9542; 1:5000), and sections were mounted in aqueous mounting medium. For vascular network analysis, fixed cardiac tissues were sectioned longitudinally at 5 µm thickness and incubated for 30 minutes at 4°C in blocking solution (10% donkey serum with 0.3% Triton X-100 in PBS). Indirect immunofluorescence was performed using goat polyclonal antibodies against CD31 (R&D Systems, AF3628; 1:30) or rat anti-CD31 (BD Pharmingen, 1:200) and α-smooth muscle actin (α-SMA; Abcam, ab32575; 1:500), followed by Cy3-and Cy5-conjugated secondary antibodies. Serum amyloid A3 (SAA3) immunostaining was performed on 5 µm cardiac sections after antigen retrieval with 70% formic acid (Thermo Scientific, #270480010) for 20 minutes at room temperature. Sections were then incubated with mouse monoclonal anti-SAA (Abbexa, abx018140; 1:50) for 2 hours at 37°C, followed by Alexa Fluor 488-conjugated anti-mouse secondary antibody. Nuclei were counterstained with DAPI and mounted using Vectashield Vibrance antifade medium (H-1000). To assess cardiomyocyte cross-sectional area (CSA), sections were blocked for 30 minutes at 4°C (10% donkey serum) and permeabilized with 0.3% Triton X-100 in PBS. Cardiomyocyte membranes were stained with wheat germ agglutinin (WGA; 5 µg/mL; Thermo Fisher Scientific, W11261) as described previously (Elia, Cannavo et al. 2021), and slides were mounted with Vectashield Vibrance antifade medium (H-1000). Images were acquired with a Nikon Eclipse Ti fluorescence microscope (40× objective). For each sample, five to six random fields were analyzed (50–60 cardiomyocytes total), and diameters were measured using ImageJ software (NIH). Mean cardiomyocyte size was expressed in micrometers.

### 5.4 Cell line and experimental treatments

Human ventricular cardiomyocytes (AC16; Millipore, #SCC109; RRID: CVCL_4U18) were cultured in 10 cm² dishes and maintained at 37°C in a humidified atmosphere containing 5% CO₂. Cells were grown in DMEM/F12 medium (Lonza, Basel, Switzerland) supplemented with 12.5% fetal bovine serum (FBS) and 1% penicillin-streptomycin (P/S). Cultures were used at passages below 10 to ensure consistency and viability. AC16 cells were maintained at the pre-contractile developmental stage, enabling reliable observation of molecular responses (Davidson, Nesti et al. 2005, Lippi, Stadiotti et al. 2020), without the technical variability associated with isolated adult cardiomyocytes. Human SH-SY5Y neuroblastoma cells (ATCC, CRL-2266; RRID: #CVCL_0019, a neuronal cell line used widely in experimental neurological studies (Koriyama, Furukawa et al. 2015, Grimm, Blumel et al. 2020, Riegerova, Brejcha et al. 2021) were grown in 10 cm^2^ dishes within the 10th passage in a humidified environment at 37◦C with 5% CO_2_ in DMEM/F12 medium (Lonza Ltd. Basel, Switzerland) supplemented with 10% fetal bovine serum (FBS) and 1% penicillin-streptomycin (P/S, Lonza). For experimental treatments, equal numbers of AC16 and SH-SY5Y cells were seeded into 6-well plates. After reaching approximately 70–80% confluence (24 hours post-seeding), cells were exposed to human Aβ40 or Aβ42 oligomers (10 μM) for 16 hours. For some experiments, prior to Aβ40 exposure, cells were pretreated for 1 hour with the CAIs, ATZ (100 μM), or MTZ (300 μM) as described in previous studies (Solesio, Peixoto et al. 2018). In selected experiments, cells received a shorter 10-minute pretreatment with ATZ or MTZ, followed by a 15-minute challenge with Aβ40 oligomers (10 μM). Stock solutions of ATZ and MTZ (100 mM) were prepared in DMSO and diluted to final concentrations in serum-free medium immediately before use. Both ATZ (sc-214461) and MTZ (sc-235615) were obtained from ChemCruz. All cell cultures used in this study were routinely tested and confirmed to be free from contamination.

### 5.5 Cell and tissue homogenate preparation and quantitative immunoblotting evaluation

Cell and cardiac tissue samples were homogenized in RIPA lysis buffer (Thermo Scientific) supplemented with protease and phosphatase inhibitor cocktails (Thermo Scientific). Lysates were sonicated and centrifuged at 13,000 rpm for 10 minutes at 4°C to remove insoluble material. Protein concentration in the supernatants was determined using the Pierce BCA Protein Assay Kit (Thermo Scientific) and measured spectrophotometrically at 562 nm using the SpectraMax i3x Multi-Mode Microplate Reader (Molecular Devices). Equal amounts of protein (20–40 µg) were resolved by SDS-PAGE and transferred to membranes for western blot (WB) analysis. Total protein lysates were probed for brain-derived neurotrophic factor (BDNF; Alomone Labs, ANT-010; 1:1000), growth-associated protein 43 (GAP-43; Millipore, AB552; 1:1000), cleaved caspase-3 (Asp175; Cell Signaling, #94530; 1:500), dopamine β-hydroxylase (DβH; Millipore, AB1536; 1:1000), human amyloid-β using a combination of 4G8 (Aβ18–22; Covance, SIG-39320; 1:1000) and 6E10 (Aβ3–8; Covance, SIG-39220; 1:1000), serum amyloid A3 (SAA3, Abcam, #ab231680; 1:500), anti-rabbit-TrkB [p Tyr816] (Novus Bio, #NBP1-03499SS; 1:1000), anti-goat-TrkB (R&D Systems, AF1494; 1:1000), phospho-CREB (Ser133) (1B6) (Cell Signaling, #9196; 1:1000), CREB (Cell Signaling, #9197; 1:1000), p75NTR (Sigma Aldrich, 07-476; 1:1000), Brain Natriuretic Peptide (BNP; PA5-96084; 1:1000), and Laminin A/C (Santa Cruz Biotechnology, sc-376248; 1:500). Actin (Millipore, MAB1501; 1:1000), GAPDH (Santa Cruz Biotechnology, sc-32233, 6C5; 1:2000), and Laminin A/C were used as loading controls. Protein bands were visualized using the Odyssey® CLx Imaging System following the manufacturer’s protocol and quantified with Image Studio™ Lite Software (Marzano, Liccardo et al. 2022).

### 5.6 ELISA assay

Aβ40 and Aβ42 peptide levels were detected using a commercial ELISA kit (Invitrogen KHB3481 and KHB3441, respectively). Briefly, cardiac specimens for each mouse were collected at the end of the experimental study and homogenized with RIPA cell lysis buffer (Thermo Scientific), supplemented with phosphatases and protease inhibitors (Thermo Scientific, Waltham, MA). Then, ELISA was performed using an equal amount of total cardiac protein following the company’s specifications, and the plate was evaluated at 450 nm with a spectrophotometer reader (SpectraMax i3x Multi-mode Microplate Reader, Molecular Devices).

Cardiac mature BDNF, mature NGF, and their pro-neurotrophin isoforms (proBDNF and proNGF) were measured using commercial ELISA kits (biosensis BEK-2211/2237 and BEK-2213/2236). Cardiac tissue from each mouse was collected at study completion, homogenized in RIPA buffer (Thermo Scientific) supplemented with phosphatase and protease inhibitors, and processed according to the manufacturer’s instructions. Equal amounts of total cardiac protein were loaded for ELISA, and absorbance was measured at 450 nm using a SpectraMax i3x multimode microplate reader (Molecular Devices).

### 5.7 Preparation of Aβ oligomers

Pre-aggregated Aβ40 oligomers were prepared from hexafluoroisopropanol (HFIP)-pretreated and lyophilized peptides, following established protocols (Fossati, Giannoni et al. 2016). Briefly, lyophilized Aβ40 peptides were initially dissolved in DMSO to a concentration of 5 mM, then diluted to 100 μM in ice-cold DMEM/F12 medium. The solution was incubated at 4°C for 16 hours to promote oligomer formation. Prior to treatment, Aβ40 oligomers were further diluted to a final concentration of 10 μM in DMEM/F12 for application to cell cultures.

### 5.8 Immunofluorescence staining of SH-SY5Y neurons and AC16 cardiomyocytes

SH-SY5Y cells were fixed with 4% paraformaldehyde (PFA) for 10 minutes, followed by washing with PBS 1X and staining with a 0.1% crystal violet dye solution (Sigma Aldrich, C0775) for 15 minutes. Imaging was conducted using a Nikon NiE Fluorescence Microscope with a 4X objective, with six random areas selected from each sample for analysis. Neurite length was quantified using ImageJ software (NIH, version 1.30), with all procedures performed by a single operator in a blinded manner (Elia, Cannavo et al. 2021). Human AC16 cardiomyocytes were seeded onto sterile glass coverslips and cultured until reaching 50–60% confluence. Cells were fixed in 4% PFA for 10 minutes at room temperature, then thoroughly washed with 1× PBS. Permeabilization and blocking were performed for 30 minutes using 1× PBS containing 0.1% Triton X-100 and 2% normal donkey serum. For experimental studies, equal numbers of AC16 and SH-SY5Y cells (24 hours post-seeding) were challenged with human Aβ40 or Aβ42 oligomers (10 μM) for 16 hours. For some experiments, before Aβ40 exposure, cells were pretreated for 1 hour with the CAIs, ATZ (100 μM), or MTZ (300 μM) as described in (Solesio, Peixoto et al. 2018). In designed experiments, cells were pretreated with either ATZ (100 μM) or MTZ (300 μM) for 10 minutes, followed by a 15-minute exposure to Aβ40 oligomers (10 μM). Stock solutions of ATZ and MTZ (100 mM) were dissolved in DMSO and diluted to final concentrations in serum-free medium immediately before use. Both ATZ (sc-214461) and MTZ (sc-235615) were purchased from ChemCruz. Following preparation, AC16 were incubated with primary antibodies against brain-derived neurotrophic factor (BDNF; Abcam, ab226843; 1:250) and cAMP response element-binding protein (CREB; Cell Signaling, #9197; 1:100). To visualize the actin cytoskeleton, cells were co-stained with Alexa Fluor™ 555-conjugated phalloidin (Thermo Scientific, A34055; 1:1000). After staining, coverslips were mounted, and high-resolution images were acquired using a Nikon Eclipse Ti fluorescence microscope equipped with a 100× objective. A single-blinded operator performed all staining and image acquisition procedures to ensure consistency and eliminate bias. All cells used in the *in vitro* experiments were free of contamination.

### 5.9 Nuclear protein extraction

AC16 human cardiomyocytes were cultured in 6-well plates until confluence and subsequently treated with pre-aggregated human Aβ40 oligomers (10 μM) for 15 minutes. Before Aβ-40 oligomers treatment, cells were pretreated with either ATZ (100 μM) or MTZ (300 μM) for 1 hour. Nuclear and cytoplasmic fractions were isolated using a sequential buffer extraction method. Cells were harvested in Buffer A [10 mM HEPES (pH 7.9), 10 mM KCl, 0.1 mM EDTA, 25% NP-40, supplemented with protease inhibitors], and the lysates were transferred to 1.5 mL microcentrifuge tubes and placed on ice. Samples were centrifuged at 13,000 rpm for 10 minutes at 4°C to separate the cytosolic fraction (supernatant), which was carefully transferred to a clean tube and stored for downstream analysis. The remaining nuclear pellet was resuspended in freshly prepared Buffer B [20 mM HEPES (pH 7.9), 0.4 M NaCl, 1 mM EDTA, 10% glycerol, supplemented with protease inhibitors] and incubated on a rotator at 4°C for 2 hours to ensure complete nuclear protein extraction. Following incubation, samples were centrifuged at 13,000 rpm for 5 minutes at 4°C. The resulting supernatant, containing the nuclear protein fraction, was collected and stored at –80°C for subsequent protein quantification and immunoblotting analyses (Marzano, Liccardo et al. 2022).

### 5.10 Statistical methods

All data are presented as mean ± standard error of the mean (SEM). Statistical comparisons between two groups were performed using a two-tailed unpaired Student’s *t*-test. For comparisons involving more than two groups, one-way or two-way analysis of variance (ANOVA) was employed, as appropriate, followed by Tukey’s post hoc test to adjust for multiple comparisons. Outliers were identified and excluded using the ROUT method (Q = 1%) in GraphPad Prism (version 9.1.0, GraphPad Software, La Jolla, CA). A *p*-value ≤ 0.05 was considered statistically significant. All statistical analyses and data visualizations were conducted using GraphPad Prism version 9 (GraphPad, La Jolla, CA).

## Authors Contributions

AE and SF designed the study. AE performed and conceptualized the experiments and analyzed the data. AE wrote the paper draft and did the literature search. AC, RVT, MW, and VRG performed experiments. SF provided relevant insights, scientific supervision, critically revised the manuscript and acquired funding. SF revised the study, edited the manuscript, and offered additional literature searches.

## Acknowledgement

We acknowledge that Graphical abstract was created using BioRender.com.

## Data availability statement

The data that support the findings of this study are available from the corresponding author upon reasonable request.

---

**Supplemental Figure 1.**
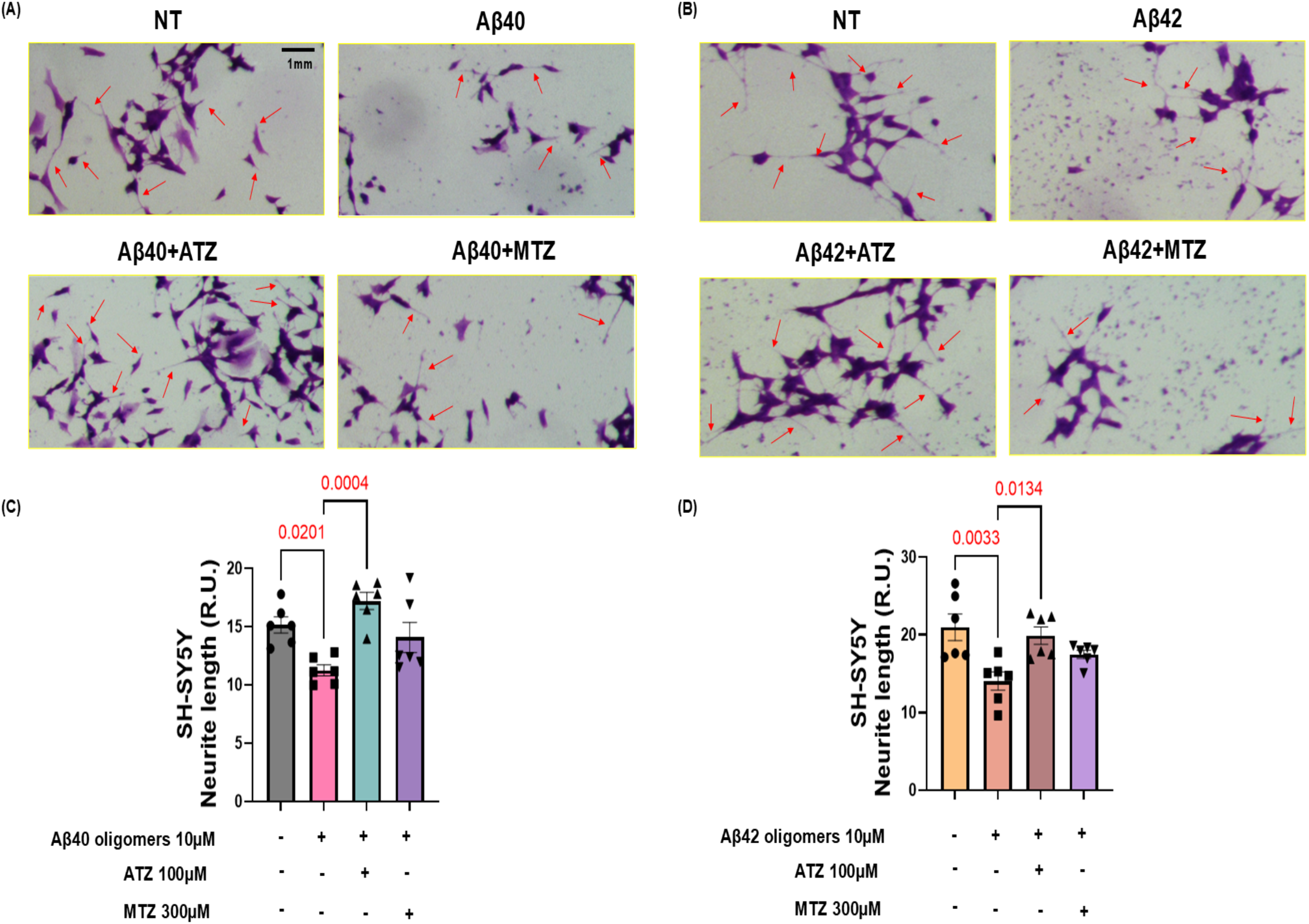
ATZ prevents Aβ-induced neurite outgrowth degeneration. (**A-B**) Representative light microscopy images of crystal violet-stained SH-SY5Y cells showing neurite morphology under non-treated (NT) control conditions or challenge with (**A**) Aβ40 or (**B**) Aβ42 oligomers (10 μM) for 16 h. Before Aβ40 or Aβ42 oligomer treatment, cells were pretreated with either ATZ (100 μM) or MTZ (300 μM) for 1 h. Red arrows indicate intact, extended neurite processes (scale bar: 1mm). (**C**-**D**) Morphometric quantification of mean neurite length per cell expressed in Relative Units (R.U.) for the (**C**) Aβ40 series and (**D**) Aβ42 series. Bars represent mean±SEM; individual data points represent replicate fields/wells. (n=6). Exact P-values indicate statistically significant differences between designated groups evaluated by one-way ANOVA. (NT = Not Treated).

**Supplemental Figure 2.**
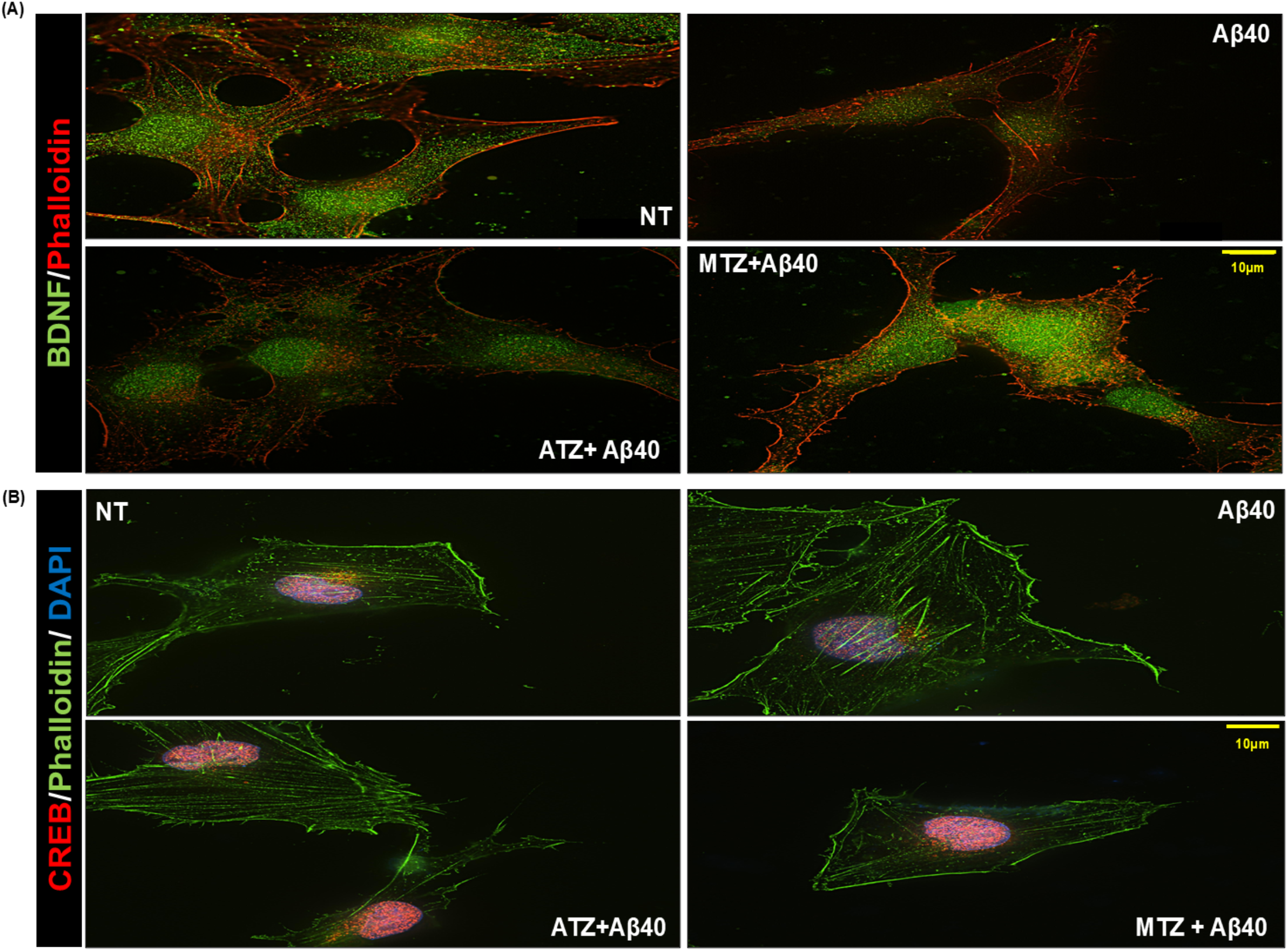
CAIs preserve intracellular BDNF and CREB signaling in Aβ40-challenged cardiomyocytes. (**A**) Representative digital images showing BDNF expression (green) and phalloidin-labeled cytoskeleton (red) in AC16 cardiomyocytes. AC16 cardiomyocytes were stimulated with Aβ40 oligomers (10 μM) for 16 h. Before Aβ40 oligomer treatment, cells were pretreated with either ATZ (100 μM) or MTZ (300 μM) for 1 h. Exposure to Aβ40 oligomers markedly reduced intracellular BDNF immunoreactivity, whereas treatment with ATZ and, particularly, MTZ, preserved neurotrophic factor expression. (**B**) Representative digital images of CREB immunostaining (red), phalloidin (green), and DAPI nuclear counterstaining (blue). AC16 cardiomyocytes were stimulated with Aβ40 oligomers (10 μM) for 15 min. Before Aβ40 oligomer treatment, cells were pretreated with either ATZ (100 μM) or MTZ (300 μM) for 10 min. Aβ40 impaired CREB signaling, while CAI treatment restored CREB expression and cellular integrity. These findings support the ability of CAIs to preserve cardiomyocyte neurotrophic signaling under amyloidogenic stress (scale bar: 10 µm). (NT = Not Treated).

**Supplemental Figure 3.**
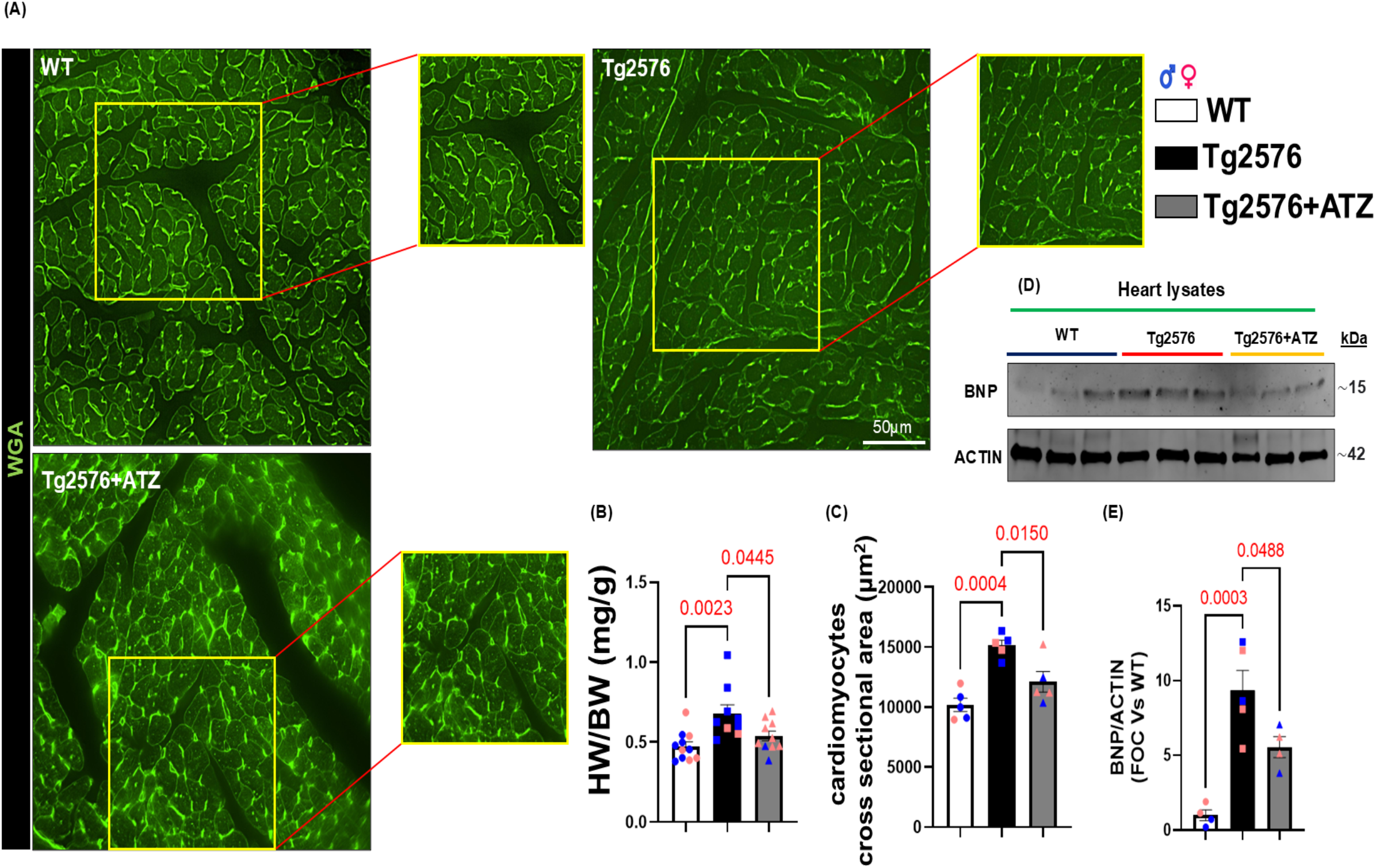
Acetazolamide treatment mitigates cardiac hypertrophy and myocardial stress in Tg2576 mice. (**A**) Representative microphotographs of ventricular tissue sections stained with Wheat Germ Agglutinin (WGA, green) to visualize cardiomyocyte membranes in 13 months old wild-type (WT), Tg2576, and Tg2576+ATZ-treated mice. Insets (red outlines) show magnified views of the quantified areas (scale bar: 50 µm). (**B**) Quantification of cardiac hypertrophy determined by heart weight/body weight (HW/BW) ratio (mg/g) across the indicated experimental groups. (WT n=11; Tg2576 n=9; Tg2576+ATZ n=10). (**C**) Histological quantification of cardiomyocyte cross-sectional area (µm^2^) derived from WGA staining (left panel). (WT n=5; Tg2576 n=5; Tg2576+ATZ n=5). (**D**-**E**) Representative Western blot (**D**) and corresponding densitometric quantification (**E**) of Brain Natriuretic Peptide (BNP) protein levels in heart lysates, normalized to ACTIN loading controls and expressed as fold over control (FOC vs WT) (right panel). (WT n=4; Tg2576 n=5; Tg2576+ATZ n=4). Individual data points are color-coded by sex (blue= male, pink= female). Data are presented as mean±SEM. Statistical significance was determined using a one-way ANOVA followed by post-hoc testing; exact p-values are indicated above the respective comparison bars.

**Supplemental Figure 4.**
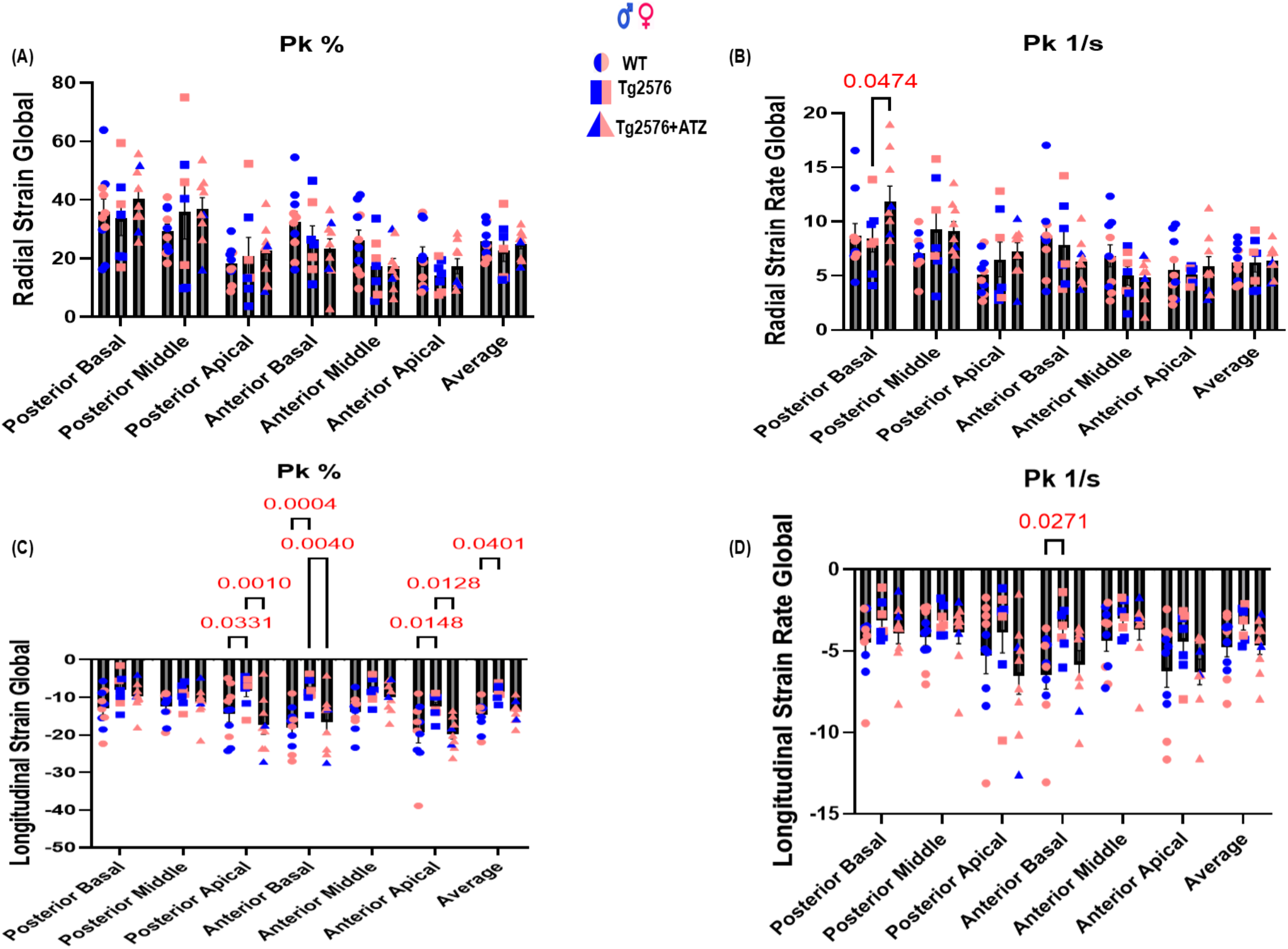
Segmental and global assessment of radial and longitudinal myocardial strain mechanics. Quantification of regional cardiac deformation and tissue velocity parameters across posterior (basal, middle, apical) and anterior (basal, middle, apical) segments, alongside the global hemispheric average, in 13 months old WT (circles), Tg2576 (squares), and Tg2576+ATZ (triangles) mice. Individual data points are color-coded by sex (blue for male, pink for female). (**A**) Peak Radial Strain Global (Pk %). (**B**) Peak Radial Strain Rate Global (Pk 1/s). (**C**) Peak Longitudinal Strain Global (Pk %). (**D**) Peak Longitudinal Strain Rate Global (Pk 1/s). (WT n=10; Tg2576 n=7; Tg2576+ATZ n=9). Data are presented as mean±SEM. Exact p-values indicate statistically significant differences based on a Two-way ANOVA with Tukey’s multiple comparisons post hoc tests (significance threshold: p < 0.05). Solid horizontal bars denote specific intergroup comparisons.

**Supplemental Figure 5.**
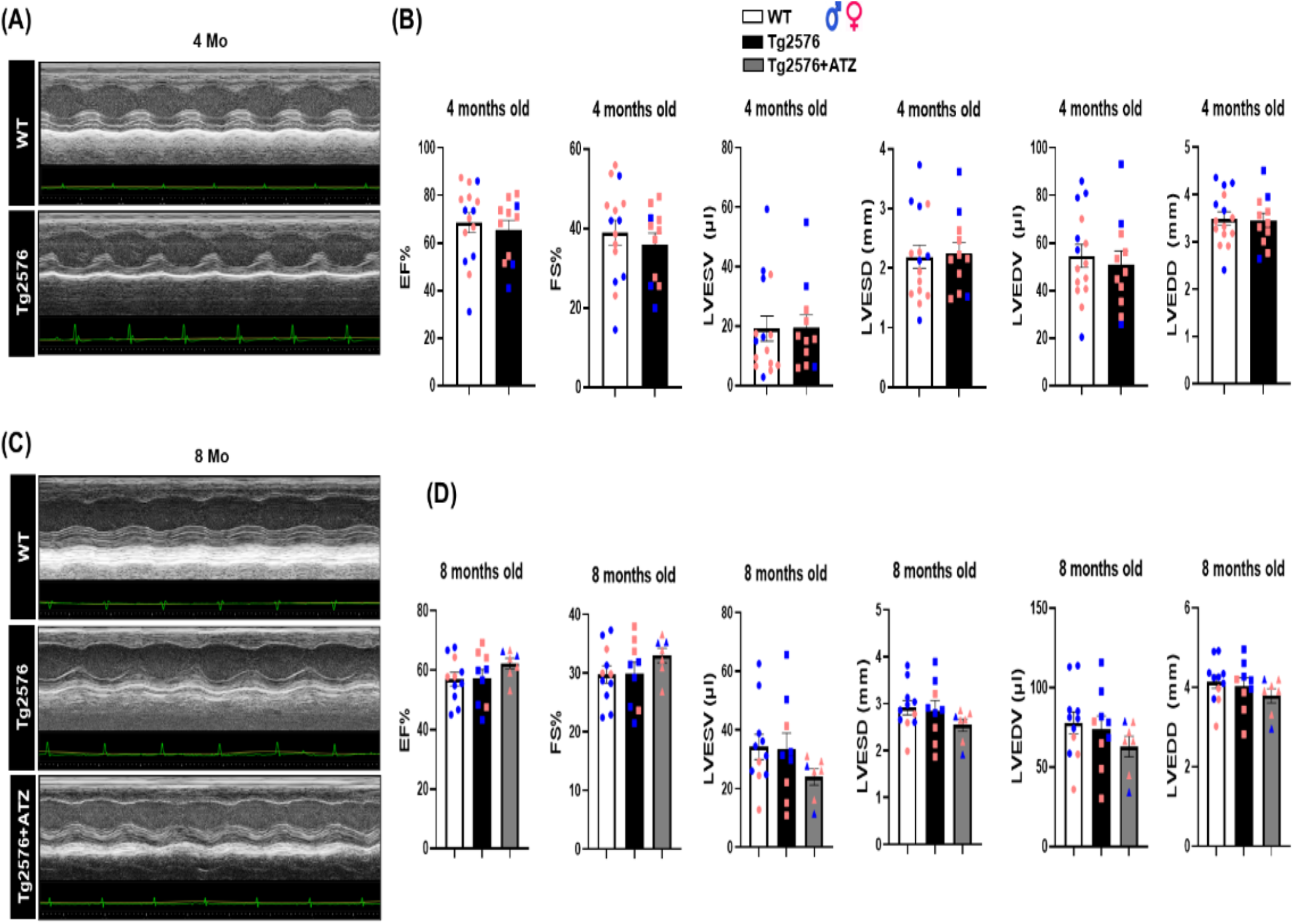
Echocardiographic assessment of cardiac structure and function at 4 months and 8 months. (**A**) Representative M-mode transthoracic echocardiography tracings of 4-month-old wild-type (WT) and Tg2576 mice. (**B**) Baseline quantification of echocardiographic parameters at 4 months of age, including ejection fraction (EF%), fractional shortening (FS%), left ventricular end-systolic volume (LVESV), left ventricular end-systolic diameter (LVESD), left ventricular end-diastolic volume (LVEDV), and left ventricular end-diastolic diameter (LVEDD). No significant differences were observed between WT and Tg2576 cohorts. (WT n=15; Tg2576 n=11). (**C**) Representative M-mode echocardiography tracings of 8-month-old WT, vehicle-treated Tg2576, and acetazolamide-treated (Tg2576+ATZ) mice. (**D**) Quantitative analysis of cardiac function and structural geometry at 8 months of age. There were no differences between WT and Tg2576 mice at this time point and ATZ administration (for 2 months) did not alter systolic function or induce adverse ventricular remodeling compared to Tg2576 or WT controls. Individual data points are color-and shape-coded to denote sex distribution (blue symbols = males; pink symbols = females). (WT n=11; Tg2576 n=9; Tg2576+ATZ n=7). Data are presented as mean±SEM; analyzed by one-way ANOVA followed by Tukey’s post hoc test (for 8-month cohorts) or unpaired Student’s t-test (for 4-month cohorts); p>0.05 across all groups.

**Supplemental Figure 6.**
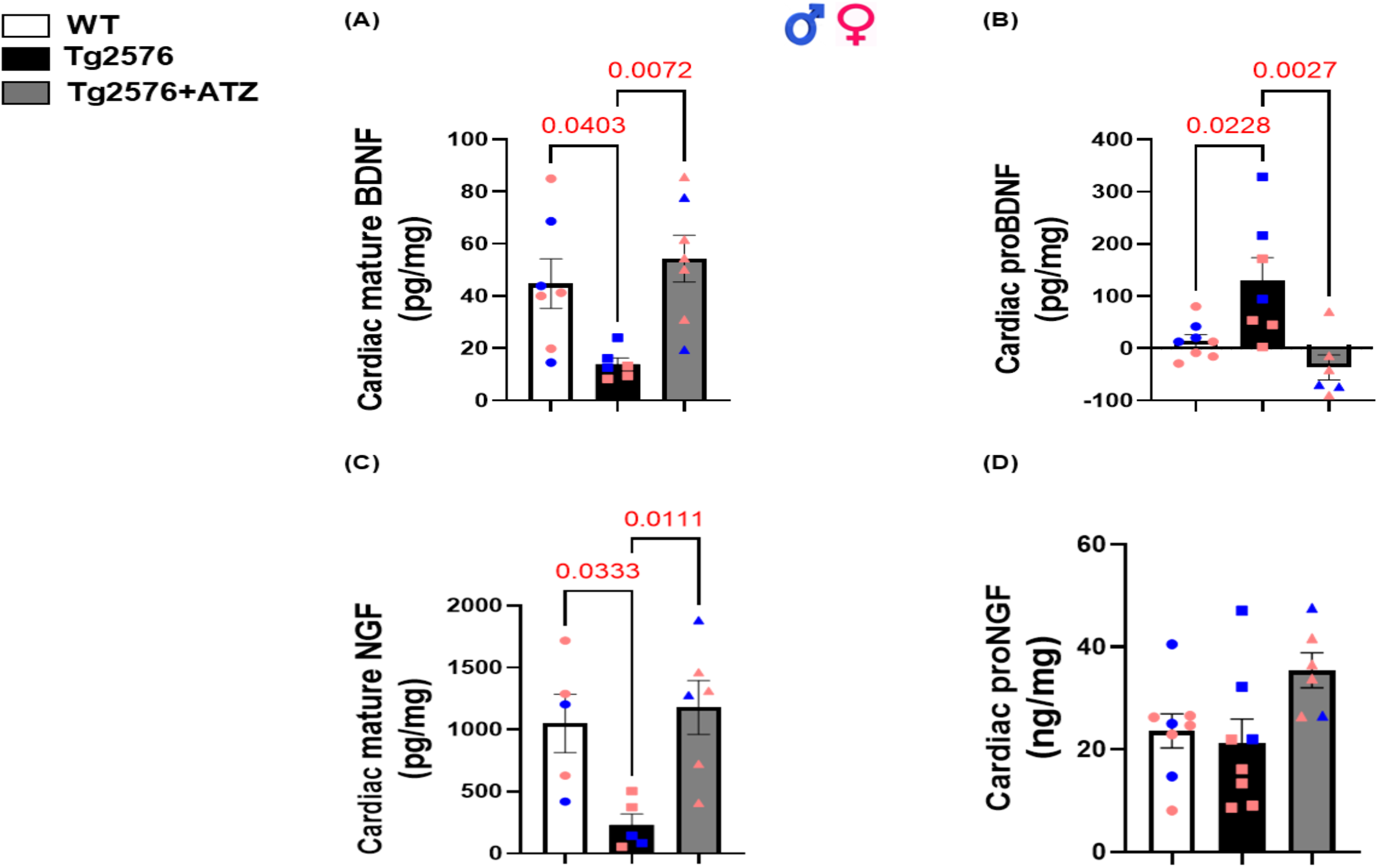
ATZ rescues cardiac neurotrophin homeostasis in 13 months old Tg2576 mice. ELISA-based quantification of (**A**) mature BDNF (WT n=7; Tg2576 n=6; Tg2576+ATZ n=7), (**B**) proBDNF (WT n=8; Tg2576 n=7; Tg2576+ATZ n=6), (**C**) mature NGF (WT n=5; Tg2576 n=5; Tg2576+ATZ n=6), and (**D**) proNGF in cardiac lysates from (WT n=8; Tg2576 n=8; Tg2576+ATZ n=7) mice. Tg2576 hearts exhibited a marked reduction in mature neurotrophins together with increased proBDNF accumulation, indicating impaired neurotrophic processing. ATZ treatment significantly preserved mature BDNF and NGF levels while reducing proBDNF accumulation, thereby maintaining myocardial neurotrophic balance. Data are presented as mean±SEM. Exact p-values are indicated above the respective comparisons, determined using a One-way ANOVA with Tukey’s post hoc test.

**Supplemental Table 1.** Echocardiographic speckle-tracking evaluation of cardiac function in WT, Tg2576, and ATZ-treated Tg2576 mice. Speckle-tracking echocardiographic parameters were assessed in 13-month-old WT, Tg2576, and Tg2576+ATZ mice: heart rate, ejection fraction (EF), radial strain and strain rate (SR) global (average of 6 LV segments), longitudinal strain and strain rate (SR) global, cardiac output; diastolic function: reverse radial and longitudinal strain global (average of 6 LV segments); dyssynchrony: radial and longitudinal strain standard deviation (STD) of T2P. (WT n=10; Tg2576 n=7; Tg2576+ATZ n=9). Data are presented as a mean±SEM. *P<0.05 vs WT, and #P<0.05 vs Tg2576+ATZ. One-way or Two-way ANOVA with Tukey’s multiple comparisons post hoc test was performed between the groups.

| Speckle-tracking Echocardiography analysis<br>*Parameters* | 13 months old |  |  |
| --- | --- | --- | --- |
|  | WT | Tg2576 | Tg2576+ATZ |
| Heart rate (bpm) | 459.84±8.18 | 484.80±11.11 | 467.87±9.27 |
| EF<sub>Strain</sub> (%) | 45.65±1.52 | 34.24±3.53**# | 43.91±1.71 |
| Radial strain Global (Pk%) | 25.76±1.84 | 22.73±3.79 | 25.02±1.65 |
| Radial SR Global (Pk1/s) | 6.24±0.52 | 6.22±0.86 | 6.43±0.46 |
| Longitudinal strain Global (Pk%) | -14.74±1.23 | -8.72±0.77**# | -13.68±0.95 |
| Longitudinal SR Global (Pk1/s) | -4.77±0.58 | -3.33±0.39 | -4.65±0.56 |
| Cardiac output (ml/min) | 14.18±1.15 | 14.24±1.18 | 11.07±0.97 |
| Reverse Radial strain global (Pk%) | -0.58±0.15 | -0.33±0.09 | -0.81±0.24 |
| Reverse Radial SR Global (Pk1/s) | -6.51±0.63 | -6.66±0.87 | -7.63±0.76 |
| Reverse Longitudinal strain global (Pk%) | 0.71±0.20 | 0.48±0.26 | 0.61±0.17 |
| Reverse Longitudinal SR Global (Pk1/s) | 4.98±0.72 | 3.19±0.32 | 4.92±0.51 |
| Radial Strain STD (T2P) | 9.01±1.06 | 10.05±1.46 | 9.50±1.04 |
| Longitudinal Strain STD (T2P) | 7.29±0.93 | 9.81±1.74 | 6.96±1.05 |

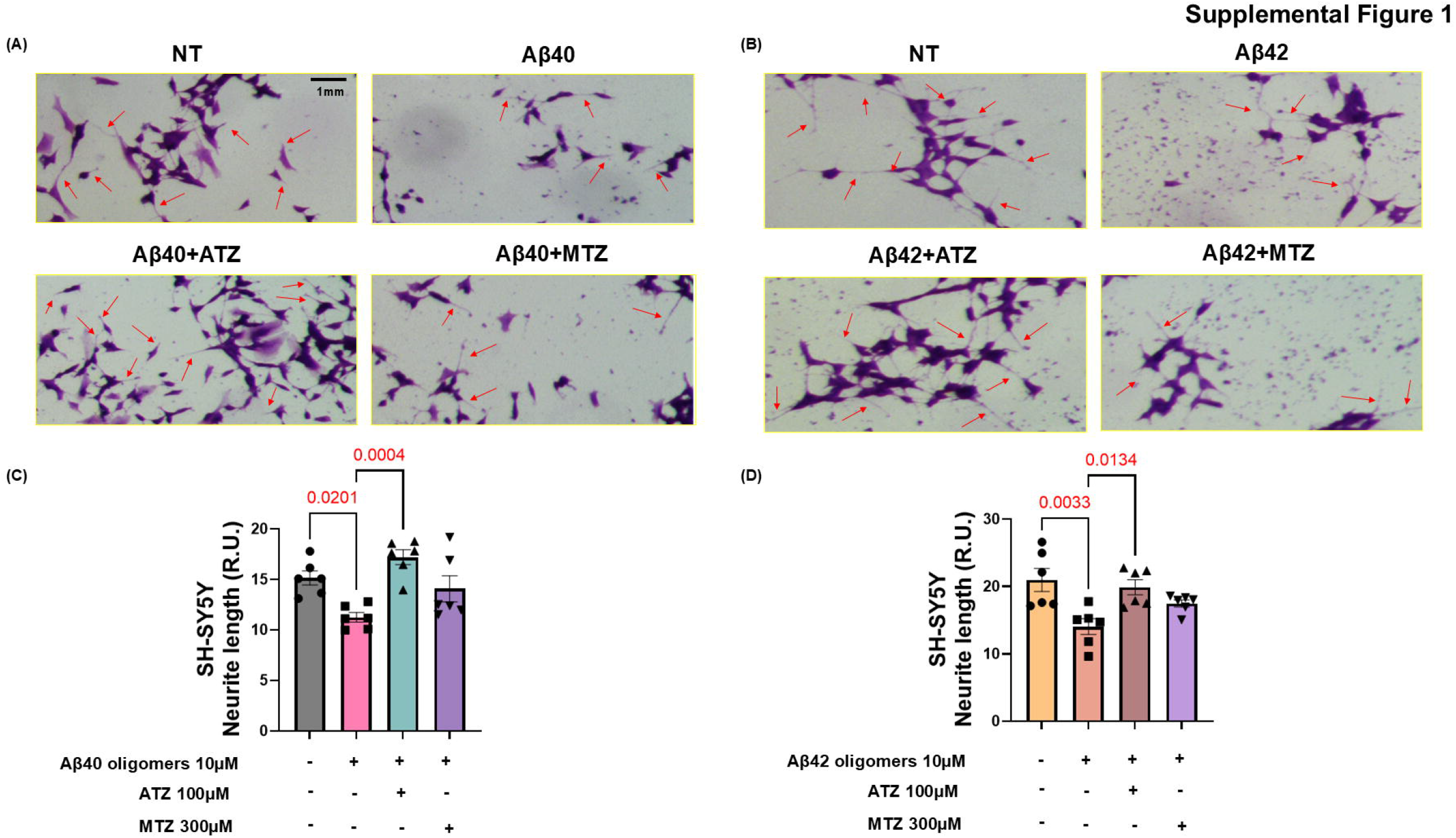

